# Deconstructing the brain bases of emotion regulation: A systems-identification approach using Bayes factors

**DOI:** 10.1101/2023.04.26.538485

**Authors:** Ke Bo, Thomas E. Kraynak, Mijin Kwon, Michael Sun, Peter J. Gianaros, Tor D. Wager

## Abstract

Cognitive reappraisal is fundamental to cognitive therapies and everyday emotion regulation. Analyses using Bayes factors and an axiomatic systems-identification approach identified four reappraisal-related components encompassing distributed neural activity patterns across two independent fMRI studies (n=182 and n=176): (1) An anterior prefrontal system selectively involved in cognitive reappraisal; (2) A fronto-parietal-insular system engaged by both reappraisal and emotion generation, demonstrating a general role in appraisal; (3) A largely subcortical system activated during negative emotion generation but unaffected by reappraisal, including amygdala, hypothalamus, and periaqueductal gray; and (4) a posterior cortical system of negative emotion-related regions down-regulated by reappraisal. These systems covaried with individual differences in reappraisal success and were differentially related to neurotransmitter binding maps, implicating cannabinoid and serotonin systems in reappraisal. These findings challenge ‘limbic’-centric models of reappraisal and provide new systems-level targets for assessing and enhancing emotion regulation.

## Introduction

Emotion regulation is a fundamental process providing contextual and goal-directed influences on feelings, decision-making, and behavior. It is crucial for everyday life and for mental and physical health (Koban et al., 2021). Dysfunctional regulation is associated with a variety of mental health and substance use disorders (Kaiser et al., 2015; Sheppes et al., 2015), as well as risk for chronic physical health conditions (DeSteno et al., 2013). One of the most effective regulatory strategies is reappraisal, which involves reinterpreting the meaning of stimuli and events in service of a regulatory goal (e.g., reducing negative emotion). Reinterpretations can include altering construals of actions (Fujita & Carnevale, 2012; Uusberg et al., 2019), causal attributions of intention (Peterson & Park, 2007), self-identification and distance, and prospections about future consequences (Gross, 2015), among other cognitive alterations (e.g. McRae et al., 2012). Regulatory goals can include suppressing or enhancing both negative and positive affect (McRae et al., 2012; McRae & Mauss, 2016; Nezlek & Kuppens, 2008). Reappraisal is a core therapeutic process in several major types of psychotherapy, such as Cognitive Behavioral Therapy (Beck, 2020; Goldin et al., 2013) and Acceptance and Commitment Therapy (Hayes et al., 2011).

A growing neuroimaging literature has examined the brain systems underlying reappraisal. Understanding the brain systems involved in emotion regulation allows them to be compared with those involved in other domain general forms of cognitive control (Ochsner & Gross, 2005), and with those affected by drugs (Baler & Volkow, 2006), brain damage, trauma, and affective processes (Heatherton & Wagner, 2011), helping to understand how such factors affect regulatory control. Identifying reappraisal-related systems also provides new neurological targets for cognitive-behavioral therapy, drug treatments, neurostimulation, and other interventions.

The major strategy for identifying reappraisal-related brain systems to date involves comparing responses to emotional stimuli with and without a regulatory goal (e.g., (Ochsner & Gross, 2005)). Regions more active during regulation are typically thought of as generators of reappraisal-related processes, whereas those with decreased activity are labeled as regulation targets. Research using this approach has found that reappraisal activates several brain regions, including the lateral prefrontal cortex, superior temporal cortex and supplementary motor area (SMA), with decreases in ‘target’ regions principally in the amygdala (Buhle et al., 2014; Kohn et al., 2014; Morawetz et al., 2017).

Several outstanding questions raised by these studies motivate the present work. The first concerns which emotion-related regions are affected by reappraisal. The amygdala is the region most consistently down-regulated by reappraisal of negative affect (Buhle et al., 2014; Frank et al., 2014), but studies and meta-analyses vary on whether amygdala activity is reduced by reappraisal (Dörfel et al., 2014). In addition, many other areas critical for generating negative affect do not appear to be consistently affected by reappraisal, including the periaqueductal gray (Etkin et al., 2015; Reddan et al., 2018), hypothalamus, ventral striatum and pallidum (Smith et al., 2009). Other cortical areas involved in affect generation also do not appear to be consistently downregulated, and some are activated during reappraisal (see below), including ventrolateral prefrontal cortex (VLPFC; (Cohen & Lieberman, 2010)), anterior insula (Craig, 2009), anterior midcingulate (Čeko et al., 2022; Shackman et al., 2011), and orbitofrontal cortex (Chikazoe et al., 2014; Salzman & Fusi, 2010). The paucity of downregulation findings could be a power issue related to the small sample sizes typical of previous studies, or could alternatively reflect a verum lack of influence, placing boundary conditions on the affective processes affected by reappraisal. In addition, affect generation is now thought to include many more regions (and distributed systems) than was appreciated in early emotion regulation studies (Barrett, 2017b; Čeko et al., 2022; Chang et al., 2015; Pessoa, 2023; Saarimäki et al., 2018), and affect generation-related regions across the brain have not been systematically identified and tested for reappraisal effects. A broader approach that identifies areas engaged during negative emotion and then assesses which of these are downregulated and which are not is needed.

Another complication is that both emotion generation and reappraisal are thought to rely heavily on cognitive appraisal (Gross & Barrett, 2011), which is important for the construction of emotion (Barrett et al., 2015) and is also linked to many of the same brain regions as those involved in reappraisal (Dixon et al., 2017). Reappraisal may thus involve many of the same appraisal processes used to generate emotions, but co-opted in service of regulatory goals (Gross & Barrett, 2011; Mesquita & Frijda, 2011). The ‘generators’ activated during reappraisal in previous studies may be specifically engaged during reappraisal, or they could be activated during negative emotion generation as well, signaling a role in common appraisal processes shared across emotion generation and regulation. Reappraisal-selective regions, should they exist, are likely associated with regulatory goals and may provide a key to understanding and assessing regulatory function in the brain. If reappraisal-selective regions can be identified, they could be measured to assess engagement in reappraisal, detect vulnerabilities, and provide brain targets for therapies or neurostimulation that are more specific to reappraisal than current brain targets.

Addressing these questions requires moving beyond traditional statistical hypothesis testing to evaluate evidence both for and against effects across multiple contrasts. For example, evaluating whether negative emotion-related regions are downregulated by reappraisal or not requires identifying emotion-relevant regions (e.g., those activated by negative vs. neutral images) and then testing evidence for and against reappraisal-related downregulation (e.g., reductions with reappraisal vs. passive viewing of negative images). Identifying reappraisal-specific regions involves establishing both a positive response to reappraisal and no difference (a null effect) when generating negative emotion in the absence of reappraisal. We refer to this as a systems identification approach, modeled on the axiomatic approach of Rutledge and Glimcher (Caplin & Dean, 2008; Rutledge et al., 2010). This approach takes the following logical form: any reappraisal-specific region must (1) respond more strongly when reappraising negative images than when merely looking at them, and (2) must not respond differentially to negative vs. neutral images in the absence of reappraisal demand. Brain regions that follow this pattern of effects can be said to track a hypothetical cognitive process that responds selectively during engagement of regulatory goals. If brain regions with these properties can be identified, support is provided that such a process is a meaningful, identifiable neural component of emotion regulation. Evidence in favor of both propositions (each involving a different contrast) can be assessed with Bayes factors, which can quantify the balance of evidence for and against the existence of an effect (Morey & Rouder, 2011). However, applying this approach has not been feasible in neuroimaging studies of reappraisal in part because establishing evidence in favor of a null effect requires substantially larger samples than have typically been available (Fu et al., 2021; Schönbrodt & Wagenmakers, 2018). For example, finding a Bayes factor with a ratio of 10:1 in favor of the null requires at least 120 participants (Supplementary Figure S2).

In this work, we included two individual fMRI datasets with large sample sizes (n=176 and n=182), focusing on findings replicated across both studies to improve generalizability across samples. Both studies scanned participants while they looked at aversive images without reappraising (‘Look Negative’), reappraised a matched set of aversive images (‘Reappraise Negative’), and looked at neutral images (‘Look Neutral’), as shown in Figure 1A. We analyzed contrasts related to emotion generation (Look Negative - Look Neutral) and reappraisal (Reappraise Negative - Look Negative). Bayes factors were calculated using a widely used, computationally efficient method (Rouder et al., 2009) that allowed efficient calculation for each brain voxel (Kragel et al., 2018). Combinations of positive and null findings were used to identify brain regions satisfying each of four potential system components (Figure 1B and 1C): (1) ‘Reappraisal only’ regions responding only to reappraisal demand, not negative images; (2) ‘Common appraisal’ regions activated by negative images and further increased during reappraisal; (3) ‘Non-modifiable emotion-generation’ regions activated by negative images but unaffected by reappraisal; and (4) ‘Modifiable emotion-generation’ regions activated by negative images and reduced by reappraisal. Figure 1B shows the profile of activity across the three task conditions for each system component, and Figure 1C shows how Bayes factors are combined across contrasts to identify voxels conforming to the axioms that define each system component. We identified regions matching each component with 10:1 evidence in favor of the required effect for each contrast (‘strong’ evidence, which also satisfies traditional FDR q < 0.05 correction for multiple comparisons in all cases), and additionally replicated the relevant effects in both studies independently. Once regions conforming to each system component were identified, we tested the correlations between averages in the relevant regions and individual differences in reappraisal success, defined as an individual’s drop in reported negative affect for [Reappraise Negative - Look Negative]. In addition, the functional and neurochemical attributes of these system components were annotated by spatial (map-wise) associations with Neurosynth topic maps derived from 11,406 previous studies (Yarkoni et al., 2011) and PET-derived molecular imaging maps from 18 neurotransmitter receptors and transporters (Hansen et al., 2022).

**Figure 1.**
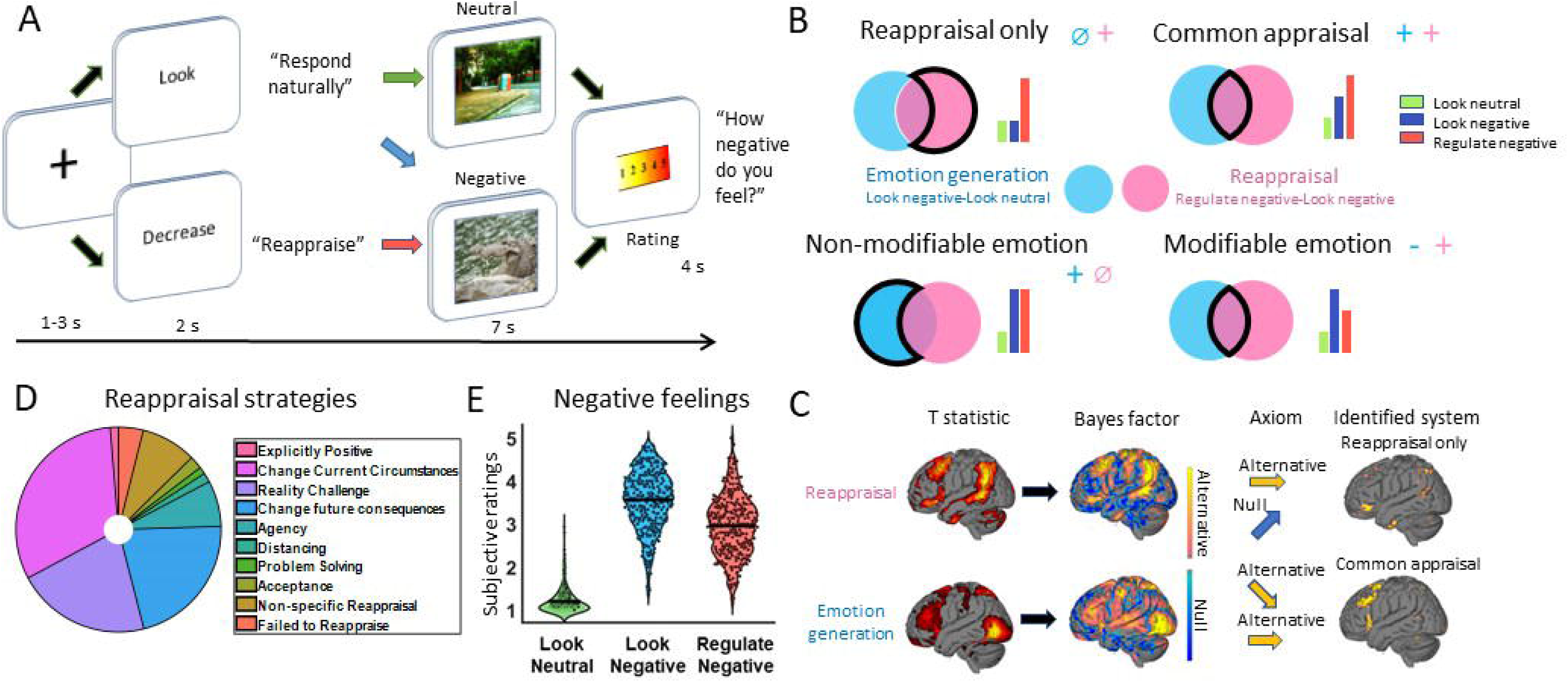
Overview of experimental paradigm and analysis pipeline. **A)** Trial structure. Sequentially, trials involved an appraisal cue (‘Look’ or ‘Decrease’, 2s), image presentation (7s), and negative affect rating (4s). On a given trial, participants either passively viewed a neutral image (‘Look neutral’), passively viewed an aversive image (‘Look negative’), or reappraised an aversive with the goal of downregulating negative affect (‘Regulate negative’). **B)** Theoretical associations between system components and brain contrasts. Circles represent Venn diagrams of voxels responding to emotion generation (Look negative vs. Look neutral, blue) and reappraisal (Regulate negative vs. Look negative, pink). ‘+’ and ‘–’ denote relative activation [+] or deactivation [-], and ⌀ denotes a null effect. The area of intersection represents an effect in both contrasts, and the non-overlapping areas are defined in part by null effects for one contrast. The portion of the Venn diagrams matching the axioms for each component are outlined in black. **C)** An example applying the axiomatic approach to identify Reappraisal-selective regions in one dataset. T-statistics for group random effects contrasts are transformed to Bayes factor maps (Rouder et al., 2009) and thresholded, with positive findings (BF>10) indicated in yellow and null findings (BF<1/10) indicated in blue. Conjunctions across thresholded Bayes factor maps identified voxels conforming to the axioms defining each system component. **D)** Specific Reappraisal Strategies. The percentage of particular reappraisal strategies used by participants. **E)** Average subjective negative ratings in each condition. Black dots represent the negative rating for individual participants averaged across trials in each condition.

## Results

### Behavioral results

Figure 1D displays the percentage of specific reappraisal strategies employed by participants in the current study, according to the coding scheme of McRae and colleagues (McRae et al., 2012). The three major strategies used by participants were ‘Change current circumstances’, ‘Reality Challenge’, and ‘Change future consequences’, indicating that strategies involving reinterpretation of the meaning of events was the major reappraisal strategy used.

Looking at negative pictures strongly increased negative affect and reappraisal decreases it on average in the current sample (Study 1+ Study 2). As shown in Figure 1E, during passive viewing (‘Look’ trials), negative affect ratings were substantially higher for negative than neutral pictures ([Look negative - Look neutral]; t = 63.8, p < 0.001, Cohen’s d = 3.37), indicating that aversive pictures increased negative affect. Reappraisal of negative pictures substantially decreased negative affect compared with passive viewing ([Regulate negative - Look negative] t = -17.5, p < 0.001, d = 0.92). 81% of individual participants showed regulation effects in the same direction as the group findings, indicating strong and robust effects.

### Emotion regulation and generation systems identified by Bayes factor analysis

How separate are neural mechanisms of reappraisal from those involved in emotion generation? We tested this question by identifying ‘Reappraisal only’ brain regions that respond to reappraisal demand (Reappraise > Look Negative, with 10:1 odds in favor of an effect) and were not activated during emotion generation (Look Negative = Look Neutral, with 10:1 odds in favor of the null). These thresholds correspond to Bayes factors (BF) of 10 and 0.1, respectively. A BF of 10 corresponds to a p-value threshold of p<0.0018, which satisfies (and is substantially more rigorous than) the q< 0.05 false discovery rate (FDR)-corrected threshold for both [Reappraise - Look Negative] and [Look negative - Look neutral]. We used the same BF thresholds in all tests reported here. In addition, we required that voxels identified in each component map (e.g., ‘Reappraisal only’) replicate all necessary effects–and thus satisfy the axioms that define the component process–in both studies independently (See “Consensus map” in Methods for details).

The consensus map for ‘Reappraisal only’ across both studies is shown in Figure 2A. We identified sixteen significant regions replicated across both datasets (Table S1). These areas were found primarily in association cortex and cerebellum, including anterior lateral prefrontal cortex (aDLPFC and aVLPFC), temporal-parietal junction (TPJ), inferior parietal lobe (IPL), posterior middle and anterior inferior temporal gyri (pMTG and aITG), and superior cerebellum. To illustrate the pattern of activation across conditions, the average responses to Reappraisal, Look Negative, and Look Neutral conditions (compared with resting baseline) in these regions are shown in the barplots in Figure 2A. To further characterize these regions, we calculated the percentage of voxels in each of the seven resting state brain networks defined by Yeo et al. (Yeo et al., 2011). Most voxels in ‘Reappraisal only’ regions fell within the default mode network (47%), frontal parietal network (31%) and ventral attention network (16%). Though nearly all regions identified in one study were replicated in the other, a small cluster in the supplementary motor area was uniquely identified in Study 1 and a small cluster in the precuneus was uniquely identified in Study 2 (Figure S1).

**Figure 2.**
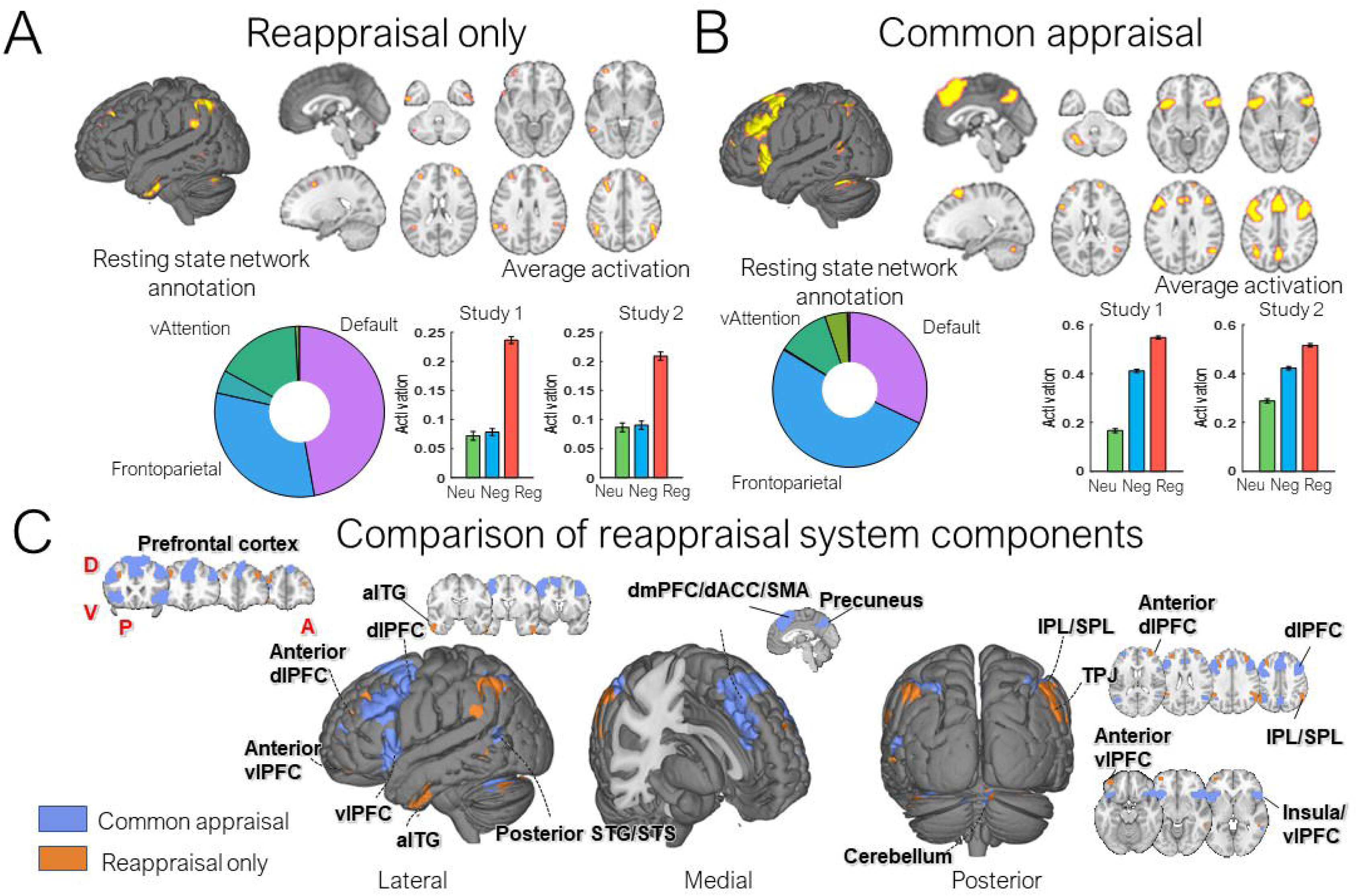
Consensus brain regions for reappraisal system components across both independent studies. (See Methods for details on consensus maps). **A)** Brain regions identified in ‘Reappraisal only’: Areas involved in emotion regulation but not generating negative emotion. Pie charts show the percentage of cortical voxels from corresponding maps located in each of the seven resting-state cortical networks from (Yeo et al., 2011). Bar plots show average data from voxels identified in each consensus map with within-subject standard error bars (Cousineau & Others, 2005). **(B)** ‘Common Appraisal’ regions: Areas positively activated in emotion generation and further increased by reappraisal. **(C)** A direct comparison of Reappraisal only (orange) and Common appraisal (blue) regions. aITG, anterior inferior temporal gyrus; dmPFC/aMCC/SMA, dorsomedial prefrontal cortex/ anterior midcingulate cortex / supplementary motor area; dlPFC, dorsolateral prefrontal cortex; vlPFC, ventrolateral prefrontal cortex; STG/STS, superior temporal gyrus/sulcus; IPL/SPL, Inferior parietal lobe/ superior parietal lobe; TPJ, temporal-parietal junction.

A motivating question for this study was whether reappraisal and emotion-generation (which involves appraisal) are represented in separate brain regions (Gross & Barrett, 2011). The results above show that some regions are specifically engaged during reappraisal. However, they do not inform on whether some regions respond to both reappraisal and emotion-generation and where such regions might be located. Accordingly, we next tested for regions that fulfill the axioms for a ‘Common appraisal’ process, which are activated in the [Look Negative - Look Neutral] contrast and further increase during reappraisal [Reappraise Negative - Look Negative].

‘Common appraisal’ regions, shown in Figure 2B, comprised the largest set of brain regions among system components identified in this study. Fourteen regions were significantly replicated across two datasets. They included dorsal and ventral lateral prefrontal cortices (DLPFC and VLPFC), superior parietal lobe (SPL), temporal sulcus (pSTS), and anterior insula (AI). In contrast to ‘Reappraisal only’ regions, ‘Common appraisal’ regions were also identified in midline brain structures, including a cluster in dorsal medial prefrontal cortex (dMPFC), anterior midcingulate cortex (aMCC), and supplementary motor area (SMA), and another cluster in precuneus. ‘Common appraisal’ regions also included cerebellum. Most voxels were located in the frontal parietal network (51%), default mode network (32%), and dorsal attention network (10%). Nucleus accumbens was uniquely identified in Study 1 but not Study 2 (Figure S1).

A closer spatial comparison of ‘Reappraisal only’ and ‘Common appraisal’ regions is shown in Figure 2C. Notably, in lateral cortical areas, areas for ‘Reappraise only’ were located anterior and lateral areas proximal to ‘Common appraisal’ regions, suggesting a cortical gradient across areas sensitive specifically to goal-directed appraisal and those spontaneously engaged by affective stimuli.

A second motivating question was to understand which regions involved in negative emotion generation are downregulated during reappraisal. We examined the set of regions activated during [Look Negative - Look Neutral], and divided them into ‘modifiable emotion’ regions reduced by reappraisal [Reappraise Negative < Look Negative] and ‘non-modifiable emotion’ regions unaffected by reappraisal [Reappraise Negative = Look Negative], using BF >= 10 in favor of the alternative or null hypothesis as above. Due to the importance of the amygdala and PAG in affect generation and prior findings of amygdala downregulation, we also tested for effects in a priori anatomically defined amygdala subregions and PAG.

Most regions responsive to emotional stimuli were non-modifiable, including all subcortical emotion-related regions (Figure 3A). Thirty-two continuous regions were identified consistently across two datasets. Non-modifiable subcortical emotion-related regions included the amygdala, periaqueductal gray (PAG), parabrachial nucleus (PBN) ventral putamen and hypothalamus, and ventral striatum and pallidum. Non-modifiable cortical regions include lateral orbitofrontal and medial prefrontal cortical regions and rectus gyrus, including midcingulate and retrosplenial cortex, agranular frontal area 6, inferior and superior parietal cortices, precuneus and ventral fusiform cortex extending into cerebellum (More details in table S1). Most cortical voxels were located in the dorsal attention network (37%) and visual system (16%) and limbic system (16%).

**Figure 3.**
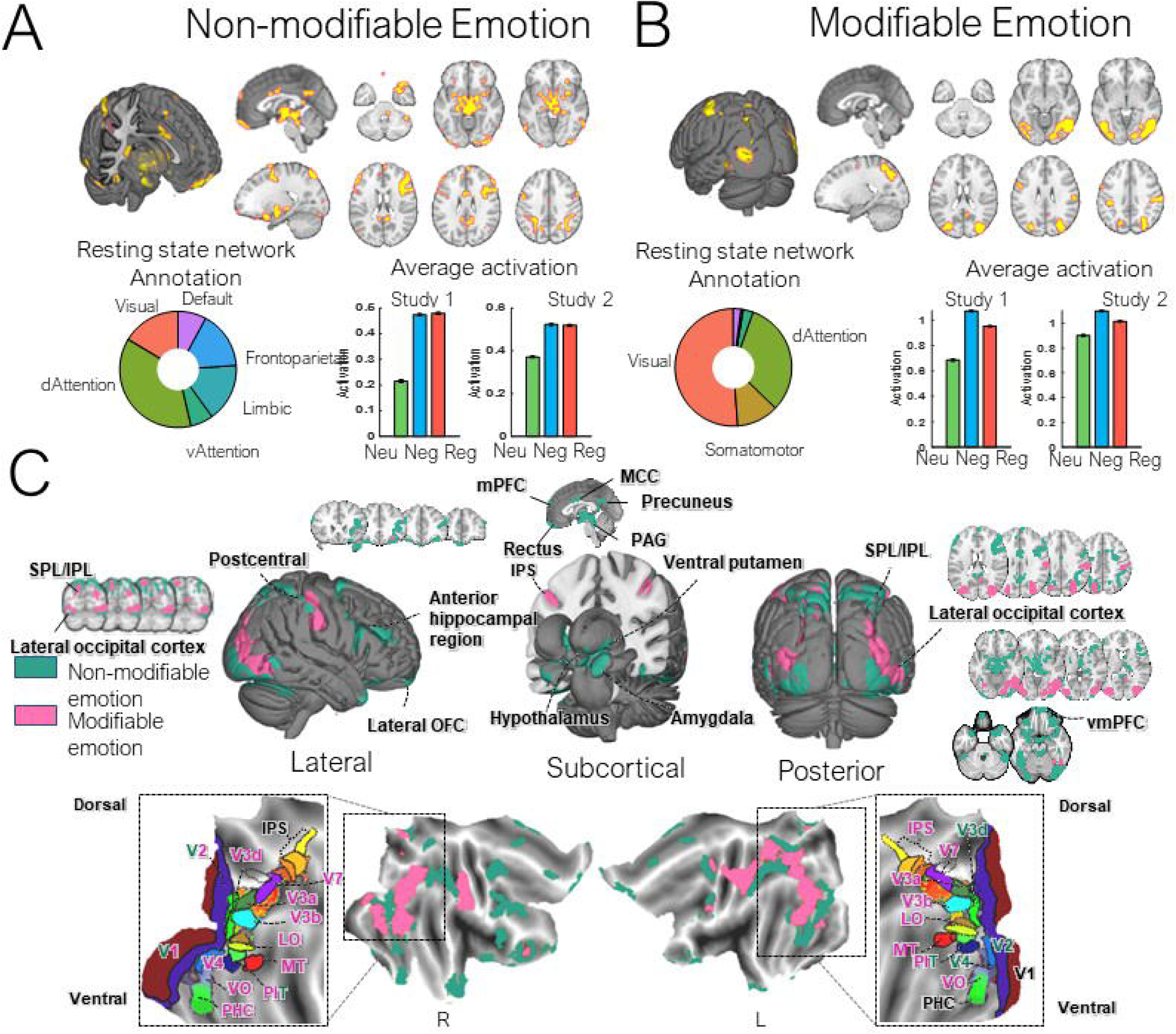
Consensus brain regions for emotion generation-related system components across both independent studies. Brain maps, pie charts, and bar plots are as in Figure 2. (A) Non-Modifiable emotion- generation brain regions: areas involved in emotion generation, but not down-regulated by reappraisal. (B) Modifiable emotion-generation brain regions: areas involved in emotion generation and down- regulated by reappraisal. (C) A direct comparison between modifiable (pink) and non-modifiable (teal) emotion-related regions. The bottom panels show regions on flattened cortical maps, with visual cortical areas labeled for reference (Van Essen et al., 2012). Modifiable and non-modifiable retinotopic visual regions are shown in pink and teal fonts, respectively, and other areas are shown in black font. IPS, intraparietal sulcus; V1, primary visual cortex; V2,V3,V4 intermediate visual cortex; MT, middle temporal; LO, lateral occipital; PIT, posterior inferior temporal (Wang et al., 2015); VO, ventral occipital; PHC, parahippocampal cortex. Abbreviation for other regions: MCC, midcingulate cortex; PAG, periaqueductal gray; OFC, orbitofrontal cortex; mPFC, medial prefrontal cortex.

‘Modifiable emotion-generation regions’ were located exclusively in the cortex, and mainly in occipito-temporal and parietal regions. Eight continuous regions were identified consistently across two datasets. The two largest clusters were located in the bilateral middle/inferior occipital cortex, overlapping with early visual regions (including the lateral occipital complex). Other regions included the postcentral gyrus, overlapping with the primary somatosensory cortex, intraparietal cortex, and inferior frontal junction in posterior prefrontal cortex. Voxels were mostly located in the three resting-state networks: the visual network (51%), dorsal attention network (31%) and somatomotor network (12%). Interestingly, ventral medial prefrontal and medial orbital prefrontal cortices were identified as modifiable only in Study 1, and a more dorsal medial prefrontal region was modifiable only in Study 2 (see Supplementary FigureS1).

A direct comparison between ‘Modifiable’ and ‘Non-modifiable’ cortical emotion-generation regions (Figure 3C), showed that both of them were located in posterior cortices. Specifically, the ‘Modifiable’ regions were mainly found in retinotopic visual areas such as the dorsal visuospatial cortex (V7) and intraparietal sulcus (IPS), ventral visual stream (V3, V4, VO and PHC), and lateral visual cortex (LO, MST, and MT), while the occipital part of the ‘Non-modifiable’ regions was mainly located in non-retinotopic areas. In the parietal lobe, most of the emotion- generation regions in the superior parietal lobe were non-modifiable, whereas the precuneus and posterior IPL were modifiable. Notably, a large subcortical emotion cluster was identified as non-modifiable, while we did not detect any modifiable emotion regions in the subcortical cortex.

Finally, we also observed ‘Non-modifiable emotion-generation’ regions in frontal regions, indicating the presence of emotion-generation specific structures in the frontal cortex as well.

ROI analyses on amygdala and PAG regions are shown in Figure 4. All amygdala subregions defined based on the anatomical tracings of Amunts et al. (Amunts et al., 2020) (implemented in the SPM Anatomy Toolbox (Laird et al., 2013) and the PAG responded robustly to negative images ([Look negative - look neutral], all p < 0.0001), indicating sensitivity to negative images in this sample. However, no region was significantly downregulated by reappraisal ([Regulate negative - look negative], all BF < 0.1), and effects in the basolateral and superficial amygdala were numerically in the opposite direction as predicted by the downregulation hypothesis.

**Figure 4.**
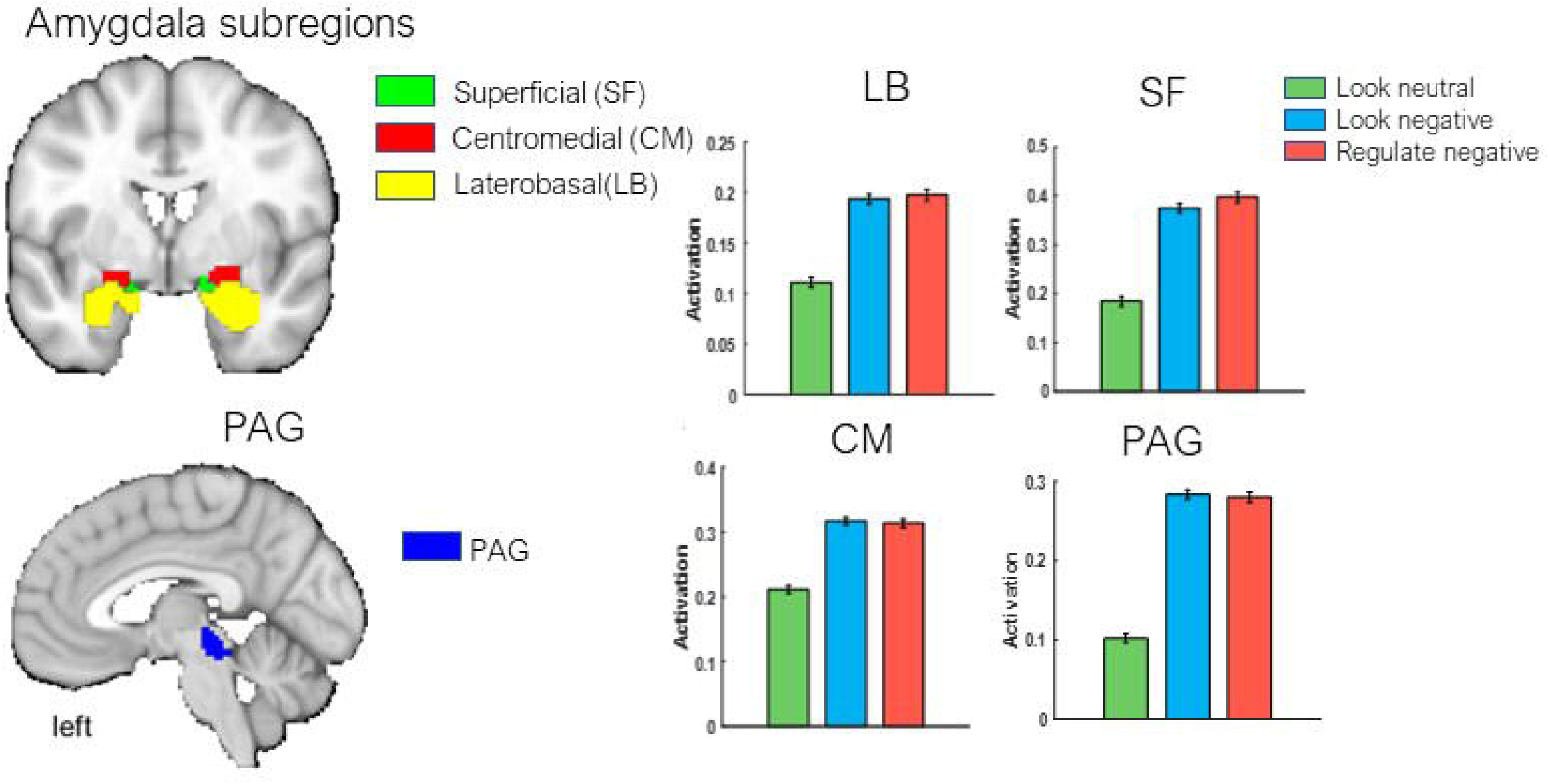
ROI analysis on amygdala and PAG. Amygdala regions are defined based on (Amunts et al., 2020), and the PAG region was defined anatomically by T.D.W. in 2018. All regions are available in the Cognitive and Affective Neuroscience Lab 2018 combined atlas (see canlab.github.io). The brain activations for each condition are shown in the bar plot.

### Covariation of reappraisal success with identified system components

In both datasets, reappraisal successfully reduced negative experience ratings. We tested whether reappraisal-related activity in the identified system components was associated with individual differences in reappraisal success (Figure 5A), defined as the average experience rating difference for [Look Negative - Reappraise Negative]. More positive scores indicated greater downregulation success. Success was correlated with increased average activity in ‘Reappraisal only’ (r=0.2, p=0.0002, adjusted R square=0.039) and ‘Common appraisal’ (r=0.14, p=0.008, adjusted R square=0.02) systems, decreased activity in the ‘Modifiable emotion-generation’ system (r=-0.14, p=0.008, adjusted R square=0.017). These correlation values are similar in strength to the strongest univariate associations across many phenotypes tested in recent comprehensive analyses (Marek et al., 2022). Average activity in the ‘Non-modifiable emotion-generation’ system was not significantly correlated with reappraisal success (r=0.015, p=0.77, adjusted R square=-0.002).

**Figure 5.**
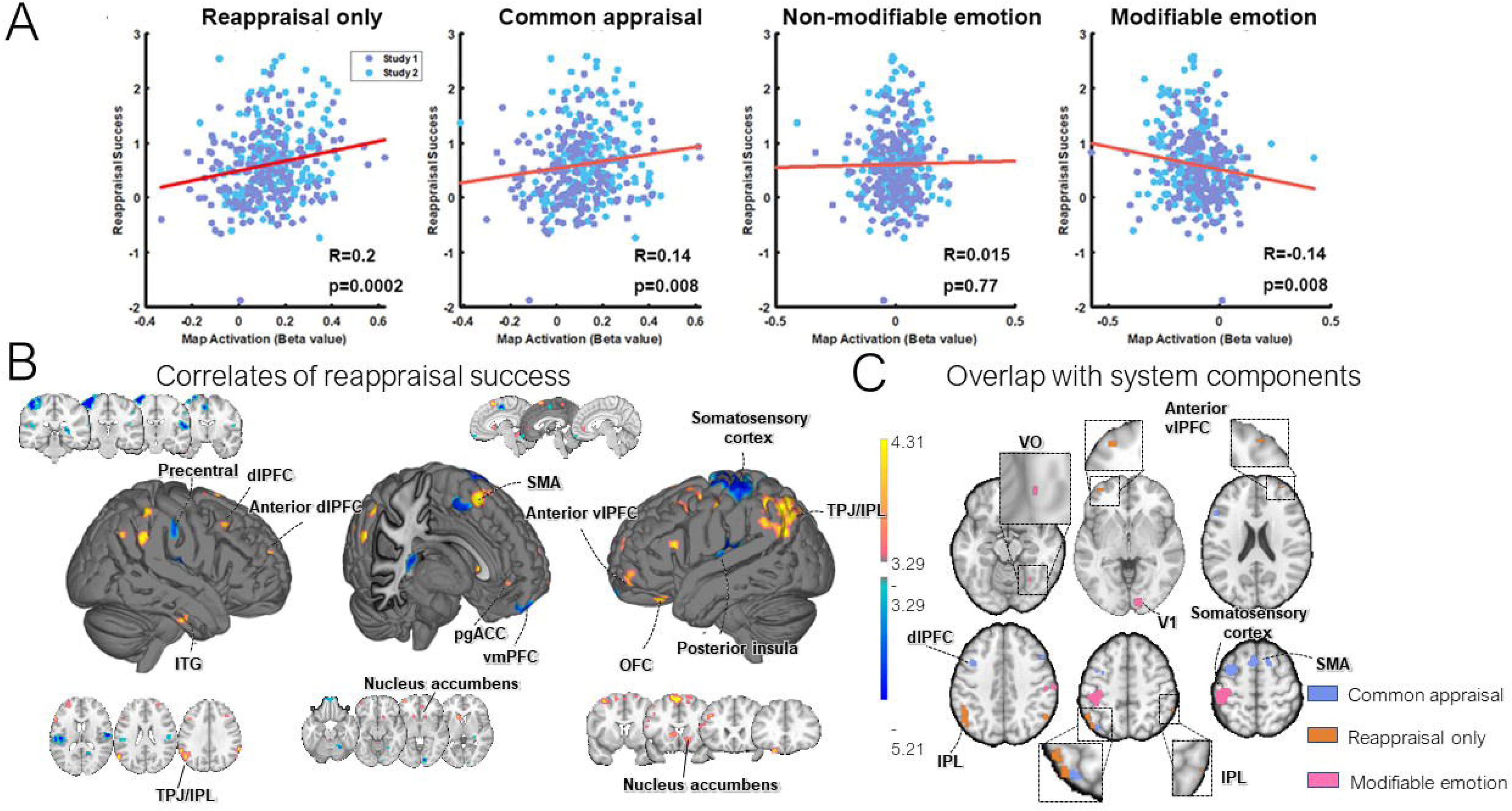
Correlation of reappraisal success with average activation from each component system. The brain activation is computed by the contrast between reappraisal-negative vs look-negative. **(A)** Scatterplots depict the cross-participant correlation between reappraisal success and average activation within each system component. Reappraisal success was correlated with increases in the average activation of ‘Reappraise only’ and ‘Common appraisal’ systems and decreases in the average activation of the ‘Modifiable emotion-generation’ system during reappraisal. Best multiple regression model to predict reappraisal success: ‘Reappraisal only’ + ‘Modifiable emotion-generation’ (Supplementary result). Regions with positive correlation show similar spatial locations with ‘Reappraisal only’ regions. **(B)** Whole brain analysis of robust regression between reappraisal success and brain activation. FDR corrected, q<0.05. **(C)** Illustrates the overlap regions between Figure 5B and ‘Reappraise only ‘, ‘Common appraisal’, and ‘Modifiable emotion-generation’ regions. The first two system components overlap with regions showing positive correlations, while the third overlaps with regions showing negative correlations. pgACC, pregenual anterior cingulate cortex.

To test whether these systems each explained unique variance in reappraisal success, and to identify the best overall model from the system component-averages, we used best subsets regression (Mallows’ CP; (Gilmour, 1996)) to examine across the four potential brain systems (see Table S2 for a summary of results across different models). The model with the lowest AIC included both ‘Modifiable emotion-generation’ and ‘Reappraisal only’ system components (r=0.26, adjust R square=0.063, p=0.000003, AIC=697). It suggests that activation of reappraisal-selective ‘generator’ regions and reductions in modifiable cortical ‘target’ regions both explain unique variance in reappraisal success, and their combination explains more than any single system component. This model provides a simple, interpretable model that can be applied to new studies.

It is possible that reappraisal success is correlated with individual differences in areas not identified in our system component analysis, including in subcortical areas of interest like the amygdala (Ochsner et al., 2004, 2012), nucleus accumbens (Wager et al., 2008), and others. To complement our systems-focused model, we examined correlates of reappraisal success across the whole brain. Previous brain correlates of reappraisal success were primarily tested in samples of n < 70 (Morawetz et al., 2016; Wager et al., 2008), but correlations do not stabilize to a 95% confidence interval of +/- 0.1 until n > 161(Schönbrodt & Perugini, 2013). With recent studies highlighting the necessity of larger samples for the estimation of stable individual differences (Marek et al., 2022; Spisak et al., 2022; Yarkoni, 2009), we sought to revisit this question with a larger sample size (n=358 across both studies). We used robust regression to minimize the influence of outliers (Wager et al., 2005) and report results at FDR q < 0.05 corrected for multiple comparisons. As shown in Figure 5B, reappraisal success was correlated with increased activation during [Reappraise Negative - Look Negative] in multiple areas within lateral prefrontal cortex (APFC, DLPFC and VLPFC), SMA, VMPFC, TPJ, and inferior temporal gyrus. These areas largely overlap with ‘Reappraisal only’ (e.g., APFC, TPJ) and ‘Common appraisal’ (e.g., SMA, DLPFC, VLPFC) system components (Figure 5C). Other regions that are not overlapped with those system components include clusters in the nucleus accumbens (NAcc) Pregenual anterior cingulate cortex (PgACC), and the superior part of the precuneus.

Conversely, reappraisal success was associated with reduced activity in several ‘Modifiable emotion-generation’ regions, including the early visual cortex (V1 area) and ventral visual cortex (VO area), and sensorimotor cortex. Other regions include S2, posterior insula and operculum. These areas were not identified as emotion- or reappraisal-related system components in our BF-based analyses because they decreased with negative emotion (Figure S3). Activation of these areas has been associated with pain and other somatic processes, and emotional feelings including social rejection (Kross et al., 2011). They were also reported in previous literature conducting similar contrast in emotion regulation (Viewing vs downregulation (Min et al., 2022)). They may be related to interoception and important for emotion regulation in addition to other system components.

Overall, these findings validate smaller-sample reports of correlations in lateral PFC, dorsal MPFC/SMA and NAcc (Modinos et al., 2010; Wager et al., 2008), and extend them by providing increased spatial precision and generalizability to a population-based sample (Gianaros et al., 2020; Hu et al., 2018).

### Functional annotation of reappraisal system component maps

An open question is how the emotion-regulation component maps we identified relate to established patterns of activity evoked by cognitive and affective studies. In particular, some prior research suggests relationships between self-regulation and cognitive control ((Casey et al., 2011; Friedman et al., 2020; Posner & Rothbart, 1998), suggesting that ‘Reappraisal only’ and ‘Common appraisal’ maps may be similar to those involved in inhibition, response selection, and other types of executive function. On the other hand, some studies suggest that prefrontal and parietal areas involved in modifying emotions may actually be distinct from those engaged by cognitive control (López-Solà et al., 2019; Wager et al., 2011; Yarkoni et al., 2011). In addition, understanding which processes are most commonly associated with modifiable areas may yield additional insights into what types of cortical processes are modified by reappraisal.

We assessed the association between each of our reappraisal-related component maps and 50 psychological topic maps from Neurosynth.org (Yarkoni et al., 2011) (Figure 6; See Table S3). The topics included all those related to mental processes–e.g face processing, inhibition, working memory, reward, motion perception, etc.--from a set of 100 topics (V4-topic-100) developed using a topic model to link activation locations and topics discussed in publications across 11,406 studies (Poldrack et al., 2012). We considered the topics most strongly correlated with each system component and those most uniquely correlated (i.e., with stronger correlations for one component than others; see Methods for details). As shown in Figure 6, the ‘Reappraisal only’ system is most strongly and selectively correlated with ‘response inhibition’, which refers to the inhibition of prepotent responses and task sets and is an type of executive function (Miyake et al., 2000). The other most strongly associated topics were ‘executive function’ and ‘empathy & interaction’. These findings suggest a close relationship between regions activated by reappraisal and those related to executive control of responses and task sets. The ‘Common appraisal’ system is most correlated with ‘working memory’, ‘task-switching’ and ‘cognitive conflict’ and unique to ‘cognitive conflict’, suggesting relationships with both executive function and value-related processes. ‘Non-modifiable emotion’ regions were most correlated with ‘emotional face’, ‘sleep & math’ and ‘working memory’ and unique to ‘sleep & math’. ‘Modifiable emotion’ regions were most strongly correlated with ‘object recognition’, ‘motion perception’ and ‘spatial attention’, and unique to ‘object recognition’. The latter findings suggest an overlap between the brain regions subserving affect and cognition, in keeping with recent neuroscience-based accounts of affect (Barrett, 2017a; Pessoa, 2023), and in particular between modification of emotion and selection of object features and properties in images. As with all such maps, the findings are hypothesis-generating, and likely reflect biases in the literature on structure-function associations.

**Figure 6.**
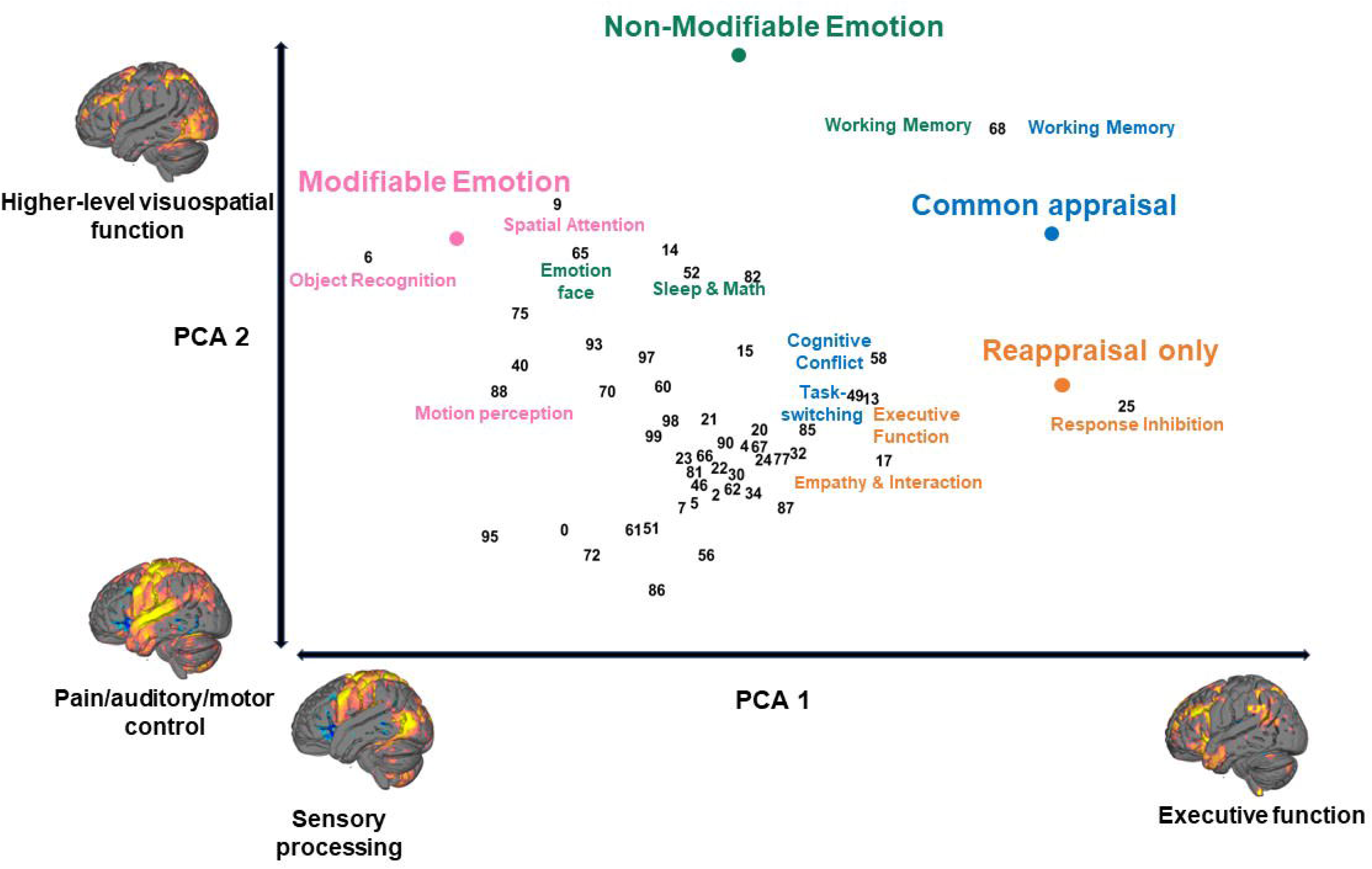
Function annotation using Neurosynth topic maps. Spatial similarities between Neurosynth topic maps and system components were assessed by Pearson correlation. This matrix is further normalized across topics within each system component map and decomposed into two dimensions using principal component analysis. All topic maps and system components were displayed on this two-dimension space given their loading on these two principal components. The brain map representing the two ends of the principal components was generated by calculating the weighted average of the top 5 loaded (negative or positive) topic maps. The top 3 correlated topics for each system component were also displayed on the graph. Additionally, we computed the most selective topic for each system component (See method for detail), which are as follows: ‘response inhibition’ for ‘Reappraisal only’; ‘cognitive conflict’ for ‘Common appraisal’; ‘emotion face’ for ‘Non-modifiable emotion’; and ‘object recognition’ for ‘Modifiable emotion’.

### Neurochemical annotation of reappraisal system component maps

It is typically unclear how fMRI-based functional systems relate to neurochemistry and neuropharmacology, as these require separate methods to directly investigate. However, recent data-sharing efforts have made it possible to relate fMRI patterns to neurotransmitter receptor and transporter binding maps obtained in Positron Emission Tomography (PET) studies (Hansen et al., 2022). In a hypothesis-generating exploratory analysis, we correlated each system component map with baseline binding maps from 36 studies of 8 different transmitter systems: cannabinoids (CB1 receptor), serotonin (5-HT receptors 1a, 1b, 2a, 4, 6, and transporter [5HTT]), acetylcholine (muscarinic M1, nicotinic a4b2, and transporter [VAChT]), norepinephrine (transporter [NET]), opioids (mu-opioid receptor), glutamate (metabotropic receptor mGluR5), histamine (H3), GABA (GABA-A, GABA-aBZ receptor), dopamine (receptors D1, D2, and transporter [DAT]). To assess error variability in the component maps, we bootstrapped the entire Bayes Factor and component-generation analysis, providing standard error bars and p-values for associations with neurochemical binding maps. This analysis does not assess error variability in the neurochemical maps themselves; therefore, we adopted a strategy of assessing replications of neurotransmitter- emotion regulation map associations across individual PET studies, which were available for many neurotransmitters (CB1, 5HT1a/1b/2a, 5HTT, VAChT, NET, mu-opioids, mGluR5, GABA, D1, D2, and DAT), and interpreted only associations that were replicated across PET studies. Agreement across PET studies appeared to be high in most cases, though there are discrepancies (note also that the same system can be assessed with different tracers, with sometimes divergent results).

As shown in Figure 7A, ‘Reappraisal only’ and ‘Common appraisal’ patterns are both significantly associated with cannabinoid (CB1), serotonin (5HT2A), mu-opioids, glutamate mGluR5, and GABA-A, suggesting the involvement of similar overall neurochemical systems (See further details in table S4). Two examples of these associations were shown in Figure 7B. The strongest correlations were with CB1 binding. 5HT1a was associated as well, but inconsistently so across PET studies for ‘Reappraisal only’, and histamine H3 binding was strongly correlated with ‘Common appraisal’ but not ‘Reappraisal only’, though only one histamine study was available so replicability could not be assessed. No other transmitter system was positively associated across PET studies.

**Figure 7.**
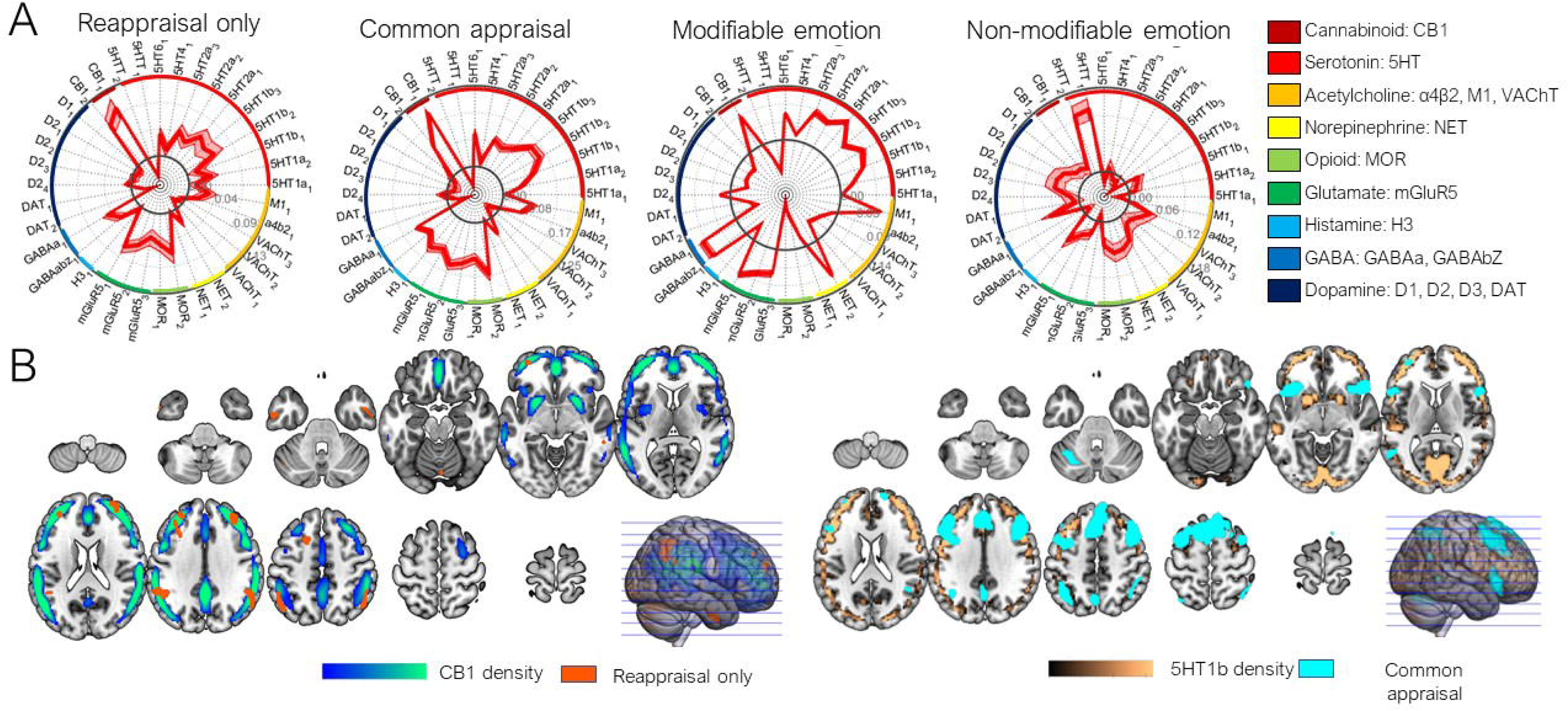
Neurochemical annotation of system component maps. **A)** Assessment of spatial similarities between individual neurochemical binding maps and system components using Pearson correlation. Error variability in component maps is evaluated through bootstrapping the entire Bayes Factor and component-generation analysis, yielding standard error bars for associations with neurochemical binding maps. **B)** Illustration of two exemplary cases of spatial overlap between neurochemical binding maps and system components.

‘Modifiable emotion’ regions were also associated with 5HT1b, 5HT2a, glutamate mGluR5, and GABA-A. The strongest numerical correlations were with GABA-A. Consistent negative correlations were found for mu-opioids, dopamine D1/D2, and VAChT, suggesting that thepresence of these neuromodulators in a region is unlikely to explain influences of reappraisal on activity. Inconsistent relationships across PET studies were found for CB1, NET, and 5HTT. In contrast, ‘Non-modifiable emotion’ regions exhibited a quite different transmitter profile, with consistent positive associations with 5HTT, NET, mu-opioids, and dopamine D2.

This pattern is consistent with fMRI findings that subcortical structures rich in dopamine, opioids, and norepinephrine were not modifiable by reappraisal.

## Discussion

Reappraisal is a foundational skill that enables success in all areas of human endeavor. However, to date, we have not had clear ways of separating brain activity related to reappraisal from activity related to the affect-generating processes it is intended to regulate. As a result, it remains unclear whether reappraisal and affect generation are similar or distinct neural processes, and we have not had brain measures to assess when and how strongly reappraisal-related neural processes are engaged. In this study, we took a step towards addressing these longstanding questions. We identified four theoretical reappraisal-related processes and found support for the existence of brain systems supporting each: (1) A reappraisal-specific process not engaged during emotion generation; (2) A common appraisal process engaged by both emotion generation and reappraisal; (3) negative emotion-related areas unaffected by reappraisal; and (4) negative emotion-related areas downregulated by reappraisal. This allowed us to localize and compare the brain regions involved in each, test how they relate to individual differences in reappraisal success (downregulation of reported negative emotion) and understand their common and distinct associations with psychological topics and neurochemical systems mapped in separate fMRI and PET studies. A key that permitted this type of analysis was the use of two large samples of over 200 participants each, which allowed us to estimate Bayes factors across the brain and quantify evidence supporting equivalent activation across conditions, as well as activation differences across conditions, with adequate power and generalizability across two independent studies (see Methods ‘Bayes factor based axiomatic method’).

### Reappraisal systems

We found that reappraisal engages distinct regions whose activity is unaffected during emotion generation alone. Reappraisal-selective areas were located anterior to ‘common appraisal’ regions in the prefrontal cortex, including aDLPFC and aVLPFC, and lateral and ventral to ‘common appraisal’ regions in the parietal cortex, temporal cortex, and cerebellum (including TPJ, aITG, and Crus I). These regions have not previously been distinguished from other regions generally activated during emotion regulation (Buhle et al., 2014; Frank et al., 2014; Kohn et al., 2014; Morawetz et al., 2017), and they may play a unique role in developing human self-regulatory capacities. TPJ, aITG, and aDLPFC are among those with the largest expansions in humans compared with non-human primates and the largest expansions during early-life human development (Hill et al., 2010).TPJ aITG, and anterior PFC also appear to be among the most integrative, transmodal areas in the brain (Margulies et al., 2016), positioned above unimodal sensory hierarchies and thus in a position to regulate them based on conceptual context. They have among the highest estimated levels of cortical hierarchy based on anatomical connections in the monkey (Burt et al., 2018) and the lowest myelin content (low myelination is a non-invasive proxy for hierarchy level (Burt et al., 2018)).

Anterior DLPFC in particular is one of the areas with the highest level of expression of human-accelerated genes (Wei et al., 2019). Functionally, aDLPFC is related to metacognitive monitoring (Shekhar & Rahnev, 2018), representation of contextual schemas (Masís-Obando et al., 2022), future-oriented thought (Okuda et al., 2003), and goal maintenance and planning (Koechlin et al., 1999). Studies of effective connectivity (Koechlin et al., 2003) and lesion studies (Badre & D’Esposito, 2009) also place it at the top-level of cortical control hierarchies. TPJ and aTC are interconnected with aDLPFC (Bludau et al., 2014) and may support similar functions. TPJ is a convergence zone for multiple cognitive processes, including attention and memory retrieval, and high-level conceptual processes including perceived agency and abstract relations (Carter & Huettel, 2013; Jung et al., 2022).

Though we identified reappraisal-selective regions, the majority of the regions activated by reappraisal were also activated during negative emotion-generation without explicit reappraisal instructions. These ‘common appraisal’ regions are presumably involved in the appraisal processes that give rise to negative emotion, and may be involved in spontaneous self-regulation in the absence of explicit regulatory instructions (Silvers et al., 2015). These regions included medial prefrontal (DMPFC, aMCC, SMA, and precuneus), lateral fronto-parietal (DLPFC, VLPFC, SPL, IPL, STS), and anterior insular cortices. These regions have been identified in previous meta-analyses of both emotion generation (Kober et al., 2008; Phan et al., 2002) and reappraisal (Kohn et al., 2014; Morawetz et al., 2017), and are co-activated in studies of emotion (Morawetz et al., 2020) (cf. MAG2 in (Morawetz et al., 2020)). These regions are located higher in connectivity-based cortical hierarchies (Burt et al., 2018) and cortical expansion over evolution and during development (Hill et al., 2010) compared to sensorimotor regions, but below reappraisal-selective regions. They span multiple resting-state networks, including ‘fronto-parietal’ and ‘default-mode’ networks, which appear to become more dynamically interconnected during affective states (Wager et al., 2015) and across multiple forms of psychopathology (Sha et al., 2019). The anterior insula in particular is thought to integrate somatic, emotional, and cognitive information to enable subjective affective awareness (Craig, 2009; Critchley et al., 2002). Other ‘common appraisal’ regions have also been associated with affective awareness, including DMPFC (Kalisch & Gerlicher, 2014; Mechias et al., 2010) and aMCC (McRae et al., 2008).

Among ‘Common appraisal’ regions, DLPFC and VLPFC are among the regions most strongly associated with self-regulation (Buhle et al., 2014), and VLPFC correlates with up-regulation of NAc and reappraisal success (Wager et al., 2008) (a finding we replicate here; see below discussion). Though they have been sometimes cast as ‘cold’ regions involved in context representation and information selection (Kohn et al., 2014; Morawetz et al., 2017; Ochsner & Gross, 2014), they may play a direct role in affective appraisal and regulation. For example, DLPFC directly correlates with trial-by-trial ratings of food craving during a self-regulation task (Hutcherson et al., 2012) and VLPFC correlates with trial-by-trial ratings of negative affect and social rejection (Chang et al., 2015; Woo et al., 2014).

Association with meta-analytic topic maps in Neurosynth show that ‘Reappraisal only’ and ‘common appraisal’ regions are involved in executive functions such as working memory and response inhibition, providing a link between appraisal and other forms of cognitive control. ‘Reappraisal only’ is involved at a less conceptual but more abstract level. Associations with neurotransmitter maps (Hansen et al., 2022; Markello et al., 2022) show that prefrontal and parietal ‘reappraisal only’ and ‘common appraisal’ regions are among the cortical regions with the highest levels of serotonin receptors (including 5HT1b and 5HT2a) and CB1 cannabinoid receptors. Serotonin systems are broadly implicated in mood, aggression, sleep, and pain. 5- HT2A is the primary receptor associated with psychedelic drug effects (Kwan et al., 2022; Vollenweider & Preller, 2020), and receptor expression is regulated by psychoactive drugs including antidepressants (Eison & Mullins, 1996). Psilocybin, a 5-HT2A agonist, has been found to reduce both negative emotions and aMCC responses associated with social rejection (Preller et al., 2015) (a ‘common appraisal’ region here). Previous studies have shown the effects of psychotherapy on serotonin binding, including 5-HT1B (Tiger et al., 2014) and 5- HT1A (Karlsson et al., 2013). CB1 is also implicated in mood regulation; e.g., genetic deficiency or CB1 antagonism can cause an increase in depression-like behavior (Shen et al., 2019). These studies, combined with our findings, suggest potential common substrates of appraisal-based and pharmacological interventions on particular molecular targets in particular brain regions.

In the context of translational science, the identification of reappraisal-selective regions and neurochemical annotations provide new targets for translational studies of psychotherapy and self-regulation training. Brain systems related to self-regulation are increasingly used as biological targets that may be engaged by psychological treatments and influenced by neurostimulation, neurofeedback, or other interventions (Gershon et al., 2003; Mayberg et al., 2005; Scheinost et al., 2013). Our findings suggest that using areas activated during reappraisal as targets will not be an effective strategy, because most areas activated by reappraisal are also activated by negative emotion-generation. Increasing activity in these areas with a neuromodulatory intervention could increase negative affect as well as potentiate reappraisal. Instead, focusing on “Reappraisal only” regions may be a better approach for these objectives. This notion is further supported by the observation that ‘reappraisal only’ demonstrates a larger and more consistent effect in predicting reappraisal success.

Neurochemical annotations are also valuable in guiding future investigations. Drugs and psychological treatments can target the same neurochemical systems and even synergize or antagonize one another (Atlas et al., 2014; Benedetti et al., 2006; Schenk et al., 2014). There are very few studies assessing the effects of psychological interventions on the various neurotransmitter systems we identify here, and a large space to explore in future studies. For example, we are aware of no attempts to assess the impact of psychotherapy or appraisal training on 5-HT2A or CB1 function, or interactions between appraisal-based and pharmacological interventions.

In summary, findings on reappraisal systems indicate that emotion generation and reappraisal depend on many of the same cortical regions, in line with psychological theories that posit overlap between affective appraisal and reappraisal. However, reappraisal involves a distinct set of brain regions that can serve as targets for assessment and intervention. These regions are situated at the highest levels of cortical hierarchies and are associated with executive cognitive functions, particularly managing hierarchies of goals over long time scales. They are also enriched in serotonin (5HT-1B and 5HT-2A), and cannabinoid (CB1) receptors, which may preferentially subserve and interact with reappraisal.

### Emotion systems

Another crucial goal of emotion regulation research is to understand which affect-generating brain processes are targeted and downregulated by reappraisal and related cognitive interventions. In previous work, amygdala has been reported as the primary target of emotion regulation (Buhle et al., 2014; Frank et al., 2014) and is frequently taken as a measure of emotion regulation effect (Goodman et al., 2014). However, the amygdala and other subcortical affect-related regions–particularly periaqueductal gray, hypothalamus, and ventral striatum and pallidum– were identified as ‘non-modifiable emotion’ regions here, and were neither influenced by reappraisal nor correlated with reappraisal success in current study. This might indicate that subcortical affective processes are less modifiable by reappraisal than previously thought.

They may be more rapidly engaged, occurring before cognitive intervention is applied (Méndez-Bértolo et al., 2016; Tamietto & de Gelder, 2010), and be more automatic and less susceptible to conceptual information (Atlas et al., 2016).

Previous studies are mixed on whether they find amygdala downregulation, and several factors could explain variations in the literature. First, amygdala downregulation may depend on the strategy used and how effectively it is deployed. A previous meta-analysis reported that one of the major strategies, reinterpretation, does not result in significant deactivation of the amygdala (Dörfel et al., 2014). The dominant strategies our participants adopted were reinterpretation- based, including altered appraisals of images’ reality (‘reality challenge’), current circumstances, and future consequences, and the agency of those depicted (Figure 1D).

Second, amygdala downregulation may vary across samples and populations. Our population here was a midlife community sample of adults from Western Pennsylvania, which was older than many previous reappraisal studies based largely on college student populations (M = 42.7 (Study 1) and 40.1 (Study 2), see Gianaros et al., 2014, 2022). Future studies could continue to examine variations across self-regulatory strategies and across populations, and should be aware of this effect and take caution when taking the amygdala as measurement when studying reappraisal or related therapy.

In contrast, the target regions most strongly downregulated by reappraisal (‘Modifiable emotion regions’) were located in the visual and somatosensory cortices. One intuitive explanation may be that participants shifted their attention from emotional to non-emotional content during reappraisal (van Reekum et al., 2007), leading to decreased activation in visual cortices during reappraisal. However, this account may not be the whole story. Previous research has demonstrated successful reappraisal even when visual attention is controlled (Bebko et al., 2014; Urry, 2010). In addition, shifting attention from emotional to non-emotional content results in decreased activity in the primary visual cortex in previous studies (Ferri et al., 2013), whereas ‘modifiable emotion’ regions are localized to higher-order visual regions (e.g., V3, V4, V7, MT, VO, MST, MT) and extended through both dorsal and ventral pathway to the temporal and parietal cortex (e.g., IPS). These regions are related to higher visual properties such as object and scene representation (Freud et al., 2016; Kravitz et al., 2013), but also encode semantic meaning and context (Huth et al., 2012). Therefore, the processes reappraisal is impacting could include semantic and conceptual representations in these areas rather than simply reflecting shifts in attended location.

Extending this view, visual and somatosensory streams are increasingly seen as integral to affect generation. Recent work suggests that human sensation and emotion are inextricably linked (Rodriguez & Kross, 2023), and predicting perceptual and action demands is an essential part of embodied views of emotion (Niedenthal, 2007). Previous studies have found that emotion triggers distinct neural representations in the visual cortex (Bo et al., 2021; Kragel et al., 2019). These neural representations are valence-specific (Bo et al., 2022; Liu et al., 2022) and different visual cortical patterns encode at least 5 distinguishable emotion categories (Bo et al., 2022; Kragel et al., 2019; Liu et al., 2022)). Other types of affect appear to be embedded in early sensory pathways as well. For example, a recent study found a superior collicular-visual pathway specific to encoding visually generated negative affect, an inferior collicular-auditory pathway specific to aurally generated negative affect, and separate somatosensory representations for thermal and mechanical pain (Čeko et al., 2022). We therefore suggest that these sensory representations of emotion may reflect our perception of the visual properties and semantic meanings of affective stimuli, as well as the construal of personal understanding about the situation implied in the images (Lieberman 2022), and that reappraisal can modify these representations. Additionally, somatosensory and posterior insular areas downregulated by reappraisal may be related to interoception (Khalsa et al., 2009). A recent study on emotion regulation also found a similar downregulation (Min et al., 2022).

Other ‘non-modifiable emotion’ regions are distributed throughout the brain, including DLPFC, OFC, IPS, premotor cortex, and early visual areas, implying that cognitive reappraisal affects only a subset of affect-generation processes. For example, some parts of DLPFC (particularly the inferior frontal junction (IFJ) and inferior frontal sulcus), DMPFC, and OFC (Brodmann’s Area 11) are engaged during negative affect but not modifiable. Some DLPFC areas that overlap ‘common appraisal’ are similar to those that predict trial-to-trial variation in affective value (Hutcherson et al., 2020). OFC regions are also among those most closely associated with appetitive and aversive value (Chikazoe et al., 2014; Morrison & Salzman, 2009; Rolls & Grabenhorst, 2008), particularly in relation to contextual variables, and in value-based reinforcement learning (O’Doherty et al., 2001; Rolls, 2019). The relative ‘immunity’ of these regions to reappraisal suggests that cognitive reappraisal may not be entirely effective in regulating the evaluation of affective stimuli. In addition, these findings update the commonly held view that prefrontal cortices are ‘cold’ cognitive structures particularly influenced by self-regulation and provide evidence for a more nuanced pattern.

Taken together, these results indicate reappraisal is able to change brain processes attributable to perceptual, semantic, and interoceptive processes related to the construction of affective value. However, it is not able to modulate subcortical-level affective experience as well as lateral prefrontal and orbitofrontal areas related to value construction.

### Limitations

Despite the successful identification of reappraisal brain systems using axiomatic methods, several methodological limitations still need to be concerned. In the axioms to define the system, we have to assume three conditions to be true: (1) brain activation from different conditions reflects different demand on appraisal processes, (2) Look negative vs look neutral can cover brain response of appraisal processes, (3) Appraisal processes do not introduce activity decreases. In other words, we are only interested in appraisal, and appraisal-related deactivation is out of scope.

These three assumed conditions could be violated. If (1) is violated, some ‘Common appraisal’ voxels identified in our study could instead be the voxels that reflect a mix of processes rather than of appraisal process. For example, a mixture of direct activation by aversive stimulus and by reappraisal-specific processes. If (2) is violated, we will miss some appraisal regions that are not activated by look negative vs look neutral. If this violation exists, some of the ‘Common appraisal’ voxels can be mislabeled as ‘Reappraisal only’. If (3) is violated, then the activity decreases should also be characterized.

To sum up, the key limitation in this study is that emotion appraisal may not be sufficiently defined by current contrast. Future studies with more delicate designs to separate emotion appraisal and other emotion responses could potentially address these issues.

## Methods

### Participants and paradigm

Data reported in this study were derived from subgroups of two studies: the Adult Health and Behavior project—Phase 2— (AHAB-2), referring to ‘Study 1’ in the main text and the Pittsburgh Imaging Project (PIP) (Gianaros et al., 2020), referring to ‘Study 2’ in the main text. Both studies recruited participants from Greater Pittsburgh area. One hundred eighty-two participants from AHAB-2 and one hundred seventy-six participants from ‘Study 2’ are included in the current study. These participants completed an emotion picture viewing and reappraisal task. Details of the task were provided in a previous study (Gianaros et al., 2014). In the task, participants viewed 30 unpleasant and 15 neutral IAPS images (Lang et al., 1997).Specifically, participants were trained and instructed to look and attend to images or decrease their negative feeling on negative images by reappraising the affective situation of pictures. Such behaviors were instructed by a 2-s cue (‘Look’ or ‘Decrease’). The images are presented after cue for seven seconds. After the picture presentation, participants rated their negative emotion state in four seconds on a 5-point scale panel (1=neutral and 5= extremely unpleasant), followed by a randomized rest period (1-3s). The entire task period comprised fifteen ‘Look neutral’ trials, fifteen ‘Look negative’ trials and fifteen’ Decrease negative’ trials. Eleven unpleasant images for ‘Decrease negative’, thirteen unpleasant images for ‘Look negative’ trials and two neutral images were overlapped across Study 1 and Study 2.

### fMRI data acquisition and preprocessing

MRI imaging data from ‘Study 1’ and ‘Study 2’ were collected on the same 3 Tesla Trio TIM whole-body scanner (Siemens, Erlangen, Germany) using a 12-channel, phased-array head coil. fMRI acquisition parameters: FOV = 205×205mm, matrix size = 64×64, TR = 2000ms, TE = 28ms, and FA = 90°.

Functional images were realigned to the first image of the series by 6-parameter rigid-body transformation, using the re-slice step to match the first image on a voxel-by-voxel basis. Before realignment, slice-timing correction was applied to IAPS fMRI task data to account for acquisition time variation in this event-related design. Realigned images were co-registered to each participant’s skull-stripped and biased-corrected MPRAGE image. Co-registered images were then normalized to Montreal Neurological Institute (MNI) space and interpolated to 2 × 2 × 2 mm voxels. Normalized images were then smoothed by a 6mm full-width-at-half-maximum (FWHM) Gaussian kernel.

### First level analysis

Within-subject level contrast is estimated by a general linear model (GLM). The task conditions (look neutral, look negative, regulate negative) were separately modeled by boxcar function and convolved with canonical HRF function (SPM 12). Within each condition, cue, picture viewing and rating period are modeled separately. The initial timing and length of these regressors are modeled according to the onset and duration of the event. High pass filter of 1/180 Hz was applied to exclude the possible low-frequency temporal drift. In addition, six head motion parameters(*x*, *y*, *z*, roll, pitch and yaw), spikes (Coded as 1 for outlier and 0 for other volumes) and time series of white matter and cerebrospinal fluid (CSF) are included as nuisance regressors. Time series of fMRI responses within standard white matter and CSF were extracted using canonical masks in CANlab tool (https://github.com/canlab/CanlabCore). Six head motion covariates were extracted from realignment parameters. Images defined as spikes if they are outside the 95% confidence regions of the cloud of images in multidimensional space. The distance in this space is referred to as mahalanobis distance. The spike number is 19±7.7 for ‘Study 2’ and 17±8.0 for ‘Study 1’, which represents approximately 5% of the scans.

Contrasts of interest included look negative vs look neutral and regulate negative vs look negative. These two contrasts were further applied for subsequent Bayes factor analysis.

### Bayes factor based axiomatic method

We established an axiomatic method based on Bayes factor to identify the proposed system components. Each process in our theoretical model must satisfy axioms—properties that must hold for any signal that encodes the component—that are defined based on the conjunction of both activation effects and null effects across two contrasts: [Look Negative – Look Neutral] (‘*Emotion Generation*’) and [Reappraise Negative – Look Negative] (‘*Reappraisal*’). Bayes Factors (BFs) is applied to quantify and compare the support evidence for both null and alternative hypotheses for each contrast. BF value reflects the likelihood ratio between alternative and null. BF values in the current study are computed using JZS prior and the method was provided in the previous study (Rouder et al., 2009). In the method, T statistics and sample size are required to compute BF value. The equation is provided in the following:

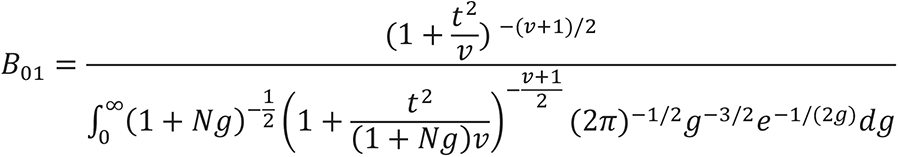

Where N stands for sample size, t stands for T statistic and v=N-1

Following the tutorial in (Lee & Wagenmakers, 2014) (Lee & Wagenmakers, 2014), we use a strict threshold of BF>10 for evidence favoring the alternative hypothesis, and BF < 1/10 favoring the null hypothesis, which is considered strong evidence. We defined system components in both datasets in following criteria:

**‘Reappraise only’** voxels have BF > 10 and t>0 for Reappraisal and BF < 1/10 for Emotion Generation. And these voxels have positive activation in ‘regulate-negative’ condition. **’Common appraisal’** voxels have BF > 10 and t>0 for both Reappraisal and Emotion Generation. And these voxels have positive activation in ‘look-negative’ condition.

**‘Non-modifiable emotion-generation’** voxels have BF > 10 Emotion Generation and BF < 1/10 for Reappraisal. And these voxels have positive activation in ‘look negative’ condition.

**‘Modifiable emotion-generation’** voxels have BF > 10 for Emotion Generation and BF > 10 for Reappraisal with a negative effect (t<0, reduced activity during reappraisal). And these voxels have positive activation in ‘look-negative’ condition.

Note the thresholds are asymmetry to null and alternative hypothesis. The null is bounded at a minimum where the means are identical (t=0). The range for T value would be narrower (BF<0.1) for null compared to strong evidence (BF>10). This range is dependent on sample size. n=120 is the minimum number to find a stable T range for null (Supplementary Figure S4). The larger sample size is better to find robust evidence for null.

### Consensus map

As shown in Figure S1, identified systems from ‘Study 1’ and ‘Study 2’ located in similar regions but not completely overlap. To identify the voxels that are spatially close to the system map from both studies, we first conducted spatial smooth using a 3mm gaussian kernel on each system component map (binary map). Next, to test the extent of overlap between ‘Study 1’ and ‘Study 2’ in each voxel, we multiplied the values in each same voxel from ‘Study 1’ and ‘Study 2’ smoothed map and yielded a product map. Higher value on the product map indicates higher degree of spatial overlap on a certain voxel. A consensus map was binary and constructed by setting a certain threshold for the product map. In current work, we use 0.01 as threshold to keep as much spatial similarity and similar number of identified voxels in consensus map compared to identified system from individual dataset.

To validate consensus maps, we tested if the voxels revealed in consensus maps overlap with or close to the identified system from ‘Study 1’ and ‘Study 2’. Specifically, we tested the distance between voxels in the consensus map and their nearest voxels in ‘Study 1’ and ‘Study 2’ systems. For each voxel, a higher value among ‘Study 1’ or ‘Study 2’ of these distances is recorded. We computed the maximum ‘Study 1’ or ‘Study 2’ distance among all voxels in the consensus map and recorded it as the ‘worst voxel’. This voxel reflects the maximum distance for the consensus map deviating from original maps. In the current parameter setting, the max deviating distance for each system component: ‘Common appraisal’, 4.4mm; ‘Reappraisal only’,4mm; ‘Non-modifiable emotion’,4.4mm; ‘Modifiable emotion’, 4.4mm.

### Neurosynth dataset

To provide an unbiased and data-driven based method to annotate the identified system components, we introduced ‘Neurosynth’ database (http://neurosynth.org), which relies on reported activation pattern from 11,406 studies (Yarkoni et al., 2011). The large scale of neurosynth dataset is able to compensate for the possible random error reported in a single study, providing robust assignment of brain function in a given region. We selected 50 related topics from a set of 100 topics (V4-topics-100) based on 11,406 studies and unrelated topics such as ‘stimulation’, ‘task’ or ‘paradigm’ are excluded. The topic maps are computed using reverse inference, which denotes probability of the topic given activation. Detailed methods to generate these topic maps can be found in Poldrack et al., 2012 (Poldrack et al., 2012). To acquire the spatial similarity between topic maps and system components, topic maps were resampled into the same space with system components. Spatial similarity is assessed via Pearson correlation between topic maps and identified systems in a pairwise fashion. Due to the large variation of voxel count across system components (‘Reappraisal only’, ‘Common appraisal’, ‘non-modifiable emotion-generation’, ‘Modifiable emotion-generation’), the scale of correlation coefficient could be biased. To balance the potential scale-level difference, we normalized the correlation coefficient within each system to make sure each SC obtains the same average correlation with 52 topics, which represents the relative correlation across all topics. A principle component analysis is conducted for the relative correlation matrix and the relative location between topics and system components are plotted on the space of first two principal components. The top 3 correlated topics for each system component were displayed.

We also reported the most unique topic for each system component compared to the other. Specifically, the uniqueness for each topic is computed by the difference between the best correlated system component and the second best one. In the end, we reported the topic that has the highest uniqueness for each system component.

### Pet-tracer neurotransmitter receptor density dataset

We investigated the spatial correlation between identified system components and a series of neurotransmitter receptor/transporter maps from a PET tracer dataset (Hansen et al., 2022). This dataset builds up whole-brain 3D normative atlases with 19 receptors and transporters across 9 different neurotransmitter systems from 1200 healthy individuals. Meta information of this dataset is shown in Table S4. Similarly, neurotransmitter receptor/transporter maps were resampled to the same space with system components for the purpose of spatial similarity.

Pearson correlation is applied to compute spatial similarity. To ensure the reliability of spatial correlation, we performed bootstrap analysis within both ‘Study 1’ and ‘Study 2’ and re- generated four system components based on 100 bootstrap samples. Bootstrap samples from each individual data are combined as a consensus map using the same method shown above. Spatial correlation is computed between bootstrapped consensus map and each PET tracer neurotransmitter map. The mean and standard deviation of spatial correlation is reported on radar plot.

## Notes

### Competing Interest Statement

The authors have declared no competing interest.

### Summary of Updates

Revise some wording in main text

