## Supplementary figure and table for "Deconstructing the brain bases of emotion regulation: A systems-identification approach using Bayes factors"

Number of Supplementary Figures: 5

Number of Supplementary Tables: 4

Figure S1: GLM contrast maps

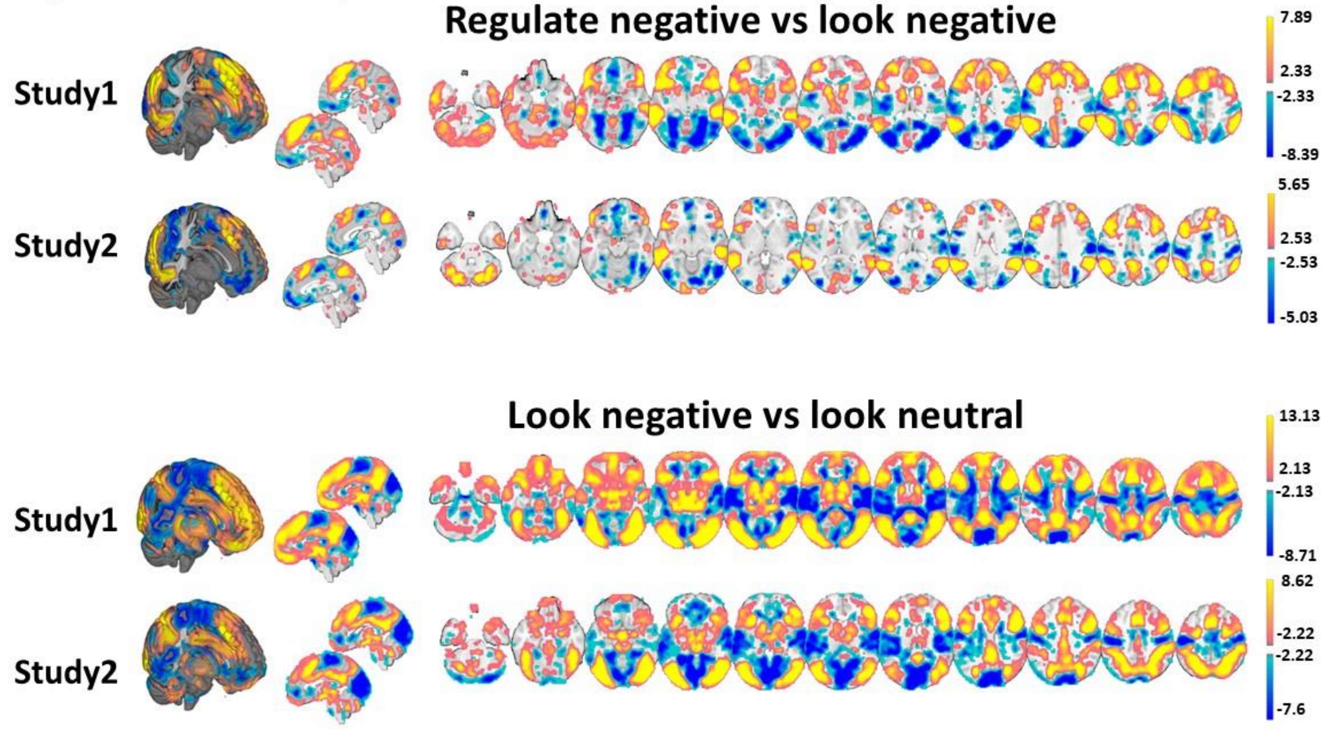


**Figure S1** 2nd-level GLM contrast T-statistic. Multiple comparisons were controlled in all maps with FDR (false discovery rate) q<0.05. Reappraisal is modeled by the contrast between 'Regulate negative' vs 'Look negative' trials. Emotion generation is modeled by the contrast between 'Look negative' vs 'Look neutral' trials. These contrast maps will be used in the subsequent Bayes factor analysis and axiomatic identification approach.

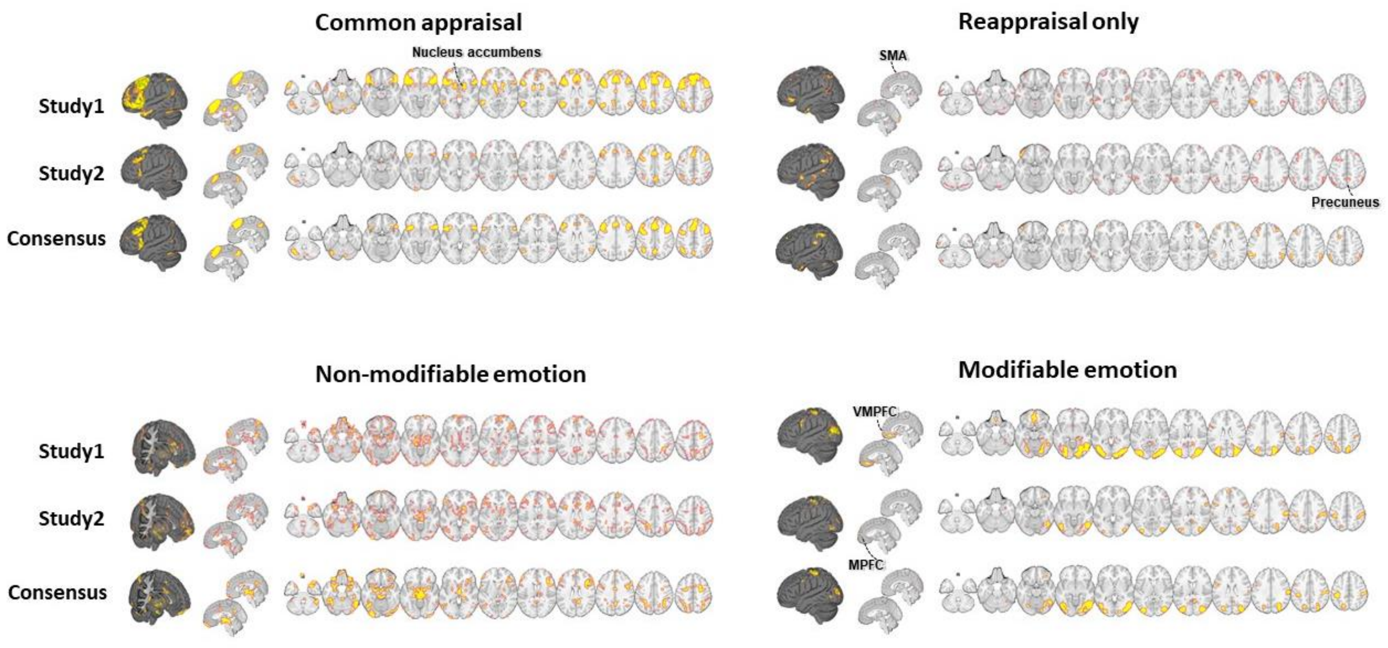


**Figure S2** Comparison of Bayes factor identified system components in each individual study and consensus map. The inconsistent regions across the two studies were labeled and corresponded to the main text.

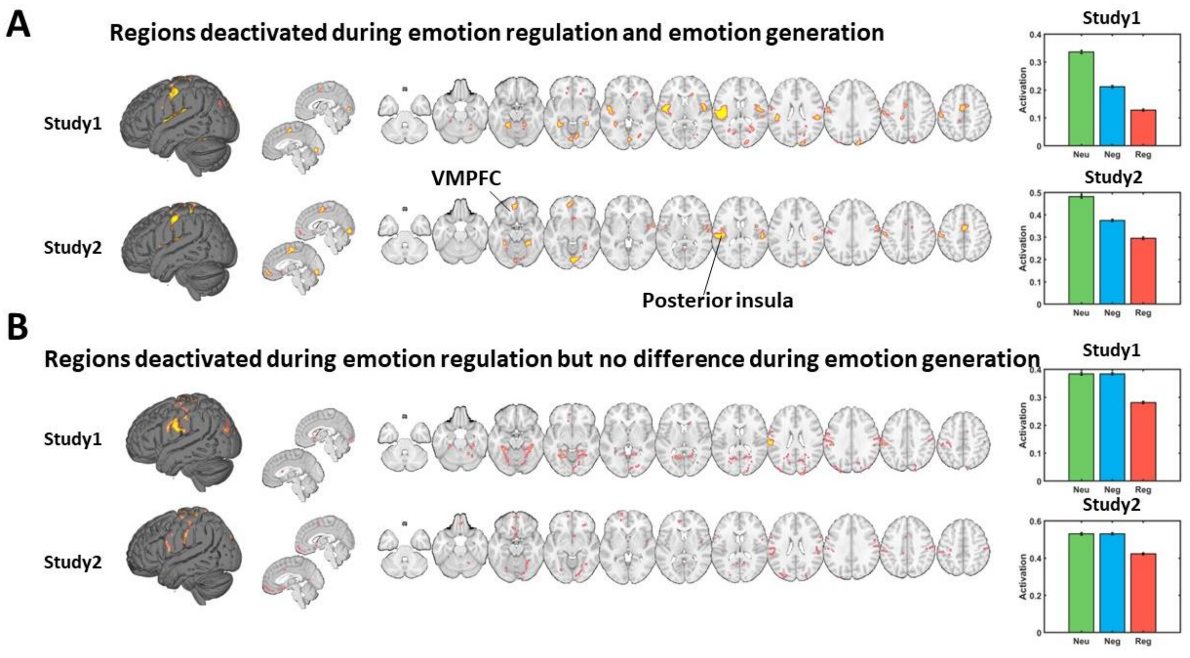


Figure S3 **Other regions deactivated during emotion regulation**
Besides 'Modifiable emotion' regions, we also found other regions that were deactivated during emotion regulation. We used the same Bayes factor axiomatic methods to identify them. **A)** Regions that were both deactivated by emotion generation and reappraisal. These regions had Bayes factors >10, t <0 for emotion generation and Bayes factors >10, t <0 for reappraisal. The average brain activations in these regions were shown on the bar plot on the right. Error bar is

computed by the same methods used in the main text. **B)** Regions that are not changed by emotion generation but deactivated by reappraisal. These regions had Bayes factors <1/10 for emotion generation and Bayes factors >10, t <0 for reappraisal. The average brain activations in these regions were shown on the bar plot on the right.

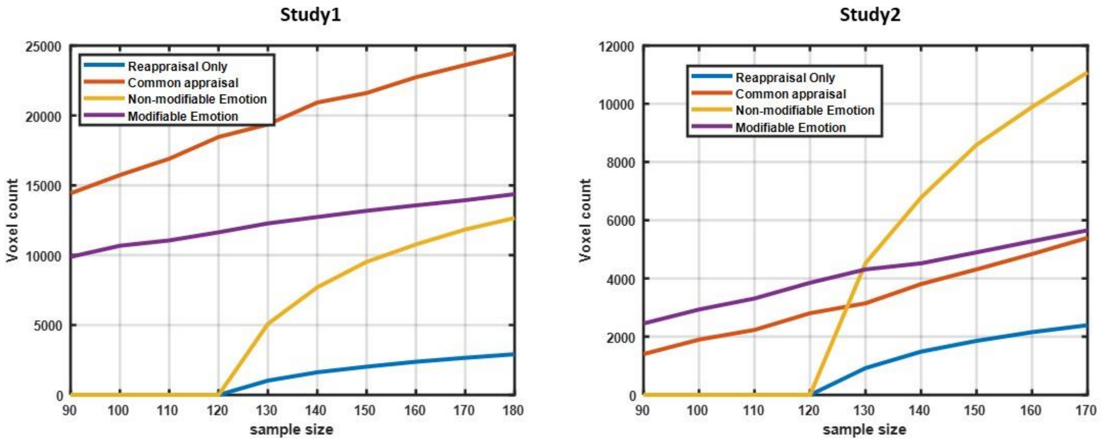


Figure S4: Assessing the impact of sample size on the detection of system components. This analysis was conducted on randomly selected subsets of each dataset, ranging from 90 to 180 participants, without repetition. This process was repeated 100 times for each subset size. The Bayes factor axiomatic approach detailed in the main text was applied to each iteration, with the number of voxels in identified systems recorded and displayed in the figure. Results indicate that reliably identifying null effects in regions such as "Reappraisal only" and "Non-modifiable emotion" necessitates a sample size of at least 120 participants, given the parameters outlined in the main text. This analysis was done for both datasets.

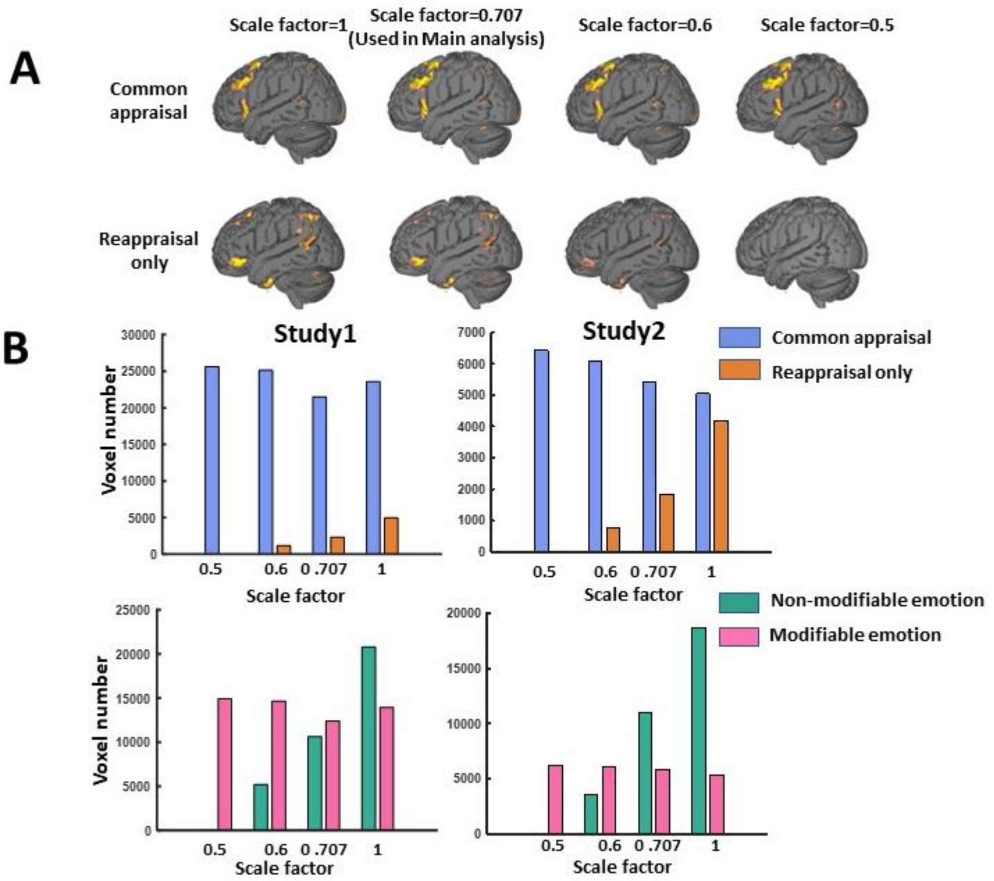


**Figure S5**: Evaluating the sensitivity of system component maps to variations in prior distribution scale factors. The scale factor represents the prior belief regarding the effect size of the data. In the main text, a scale factor of 0.707 is employed, which is a common default setting. Smaller scale factors (e.g., 0.5) may be more suitable when anticipating small effect sizes, whereas larger factors are appropriate for larger expected effect sizes. A) Visual representation of system component maps for four distinct scale factors, demonstrating that the spatial distribution of brain maps remains consistent, while the number of identified voxels varies. B) A bar plot illustrating the relationship between the number of identified voxels and scale factors. Lower scale factors result in fewer identified voxels in the "Reappraisal only" region, which requires a Bayes factor < 1/10, while smaller Bayes factors increase the number of voxels in the "Common appraisal" region, necessitating a Bayes factor > 10 for both emotion generation and reappraisal.

**Table S1** Regions that showed on figure 2 and figure 3

| | Region | CanLab_Atlas | # Voxels | Volume(mm3) | MNI Coordinates X | Y | Z | Network |
|---|---|---|---|---|---|---|---|---|
| **Common appraisal** | | | | | | | | |
| | **Cerebellum** | | | | | | | |
| L | Cerebellum | Cblm_CrusI_L | 781 | 6248 | -32 | -64 | -28 | Cerebellum |
| R | Cerebellum | Cblm_VI_R | 233 | 1864 | 10 | -74 | -26 | Cerebellum |
| | **Temporal Lobe** | | | | | | | |
| R | Superior Temporal Gyrus | Ctx_PGi_R | 340 | 2720 | 48 | -56 | 24 | Cortex_Default_ModeC |
| L | Middle Temporal Gyrus | Ctx_TPOJ2_L | 389 | 3112 | -52 | -60 | 12 | Cortex_Dorsal_AttentionA |
| R | Middle Temporal Gyrus | Ctx_PHT_R | 58 | 464 | 58 | -48 | 0 | Cortex_Dorsal_AttentionB |
| | **Parietal Lobe** | | | | | | | |
| L | Inferior Parietal Lobule | Ctx_PFm_L | 1291 | 10328 | -38 | -58 | 48 | Cortex_Fronto_ParietalB |
| R | Inferior Parietal Lobule | Ctx_PFm_R | 644 | 5152 | 44 | -54 | 52 | Cortex_Fronto_ParietalB |
| L | Precuneus | Ctx_7Pm_L | 856 | 6848 | -2 | -62 | 42 | Cortex_Fronto_ParietalC |
| | **Frontal Lobe** | | | | | | | |
| L | Middle Frontal Gyrus | Ctx_a9_46v_L | 300 | 2400 | -34 | 52 | 12 | Cortex_Ventral_AttentionB |
| R | Middle Frontal Gyrus | Multiple regions | 2553 | 20424 | 42 | 18 | 42 | Cortex_Fronto_ParietalB |
| L | Superior Medial Frontal Gyrus<br>Anterior Cingulate Cortex<br>Supplementary Motor Area | Multiple regions | 9112 | 72896 | -14 | 20 | 48 | Cortex_Fronto_ParietalB |
| R | Superior Frontal Gyrus | Ctx_9a_R | 179 | 1432 | 16 | 60 | 20 | Cortex_Default_ModeA |
| R | Insula | Ctx_44_R | 1450 | 11600 | 46 | 22 | -4 | Cortex_Default_ModeB |
| L | Insula | Multiple regions | 1526 | 12208 | -42 | 18 | -2 | Cortex_Default_ModeB |
| **Reappraisal Only** | | | | | | | | |
| | **Cerebellum** | | | | | | | |
| L | Cerebellum | Cblm_CrusI_L | 112 | 896 | -44 | -68 | -28 | Cerebellum |
| R | Vermis | Cblm_Vermis_VI | 137 | 1096 | 6 | -80 | -22 | Cerebellum |
| R | Cerebellum | Cblm_VI_R | 186 | 1488 | 28 | -70 | -26 | Cerebellum |
| | **Temporal Lobe** | | | | | | | |
| R | Inferior Temporal Gyrus | Ctx_TE1a_R | 241 | 1928 | 52 | -2 | -36 | Cortex_Default_ModeA |

| | | | | | | | | |
|---|---|---|---|---|---|---|---|---|
| L | Inferior Temporal Gyrus | Ctx_TGd_L | 303 | 2424 | -54 | -6 | -36 | Cortex_Default_ModeB |
| R | Middle Temporal Gyrus | Ctx_STSvp_R | 68 | 544 | 54 | -40 | -4 | Cortex_Default_ModeB |
| L | Middle Temporal Gyrus | Ctx_PHT_L | 104 | 832 | -58 | -52 | -2 | Cortex_Fronto_ParietalA |
| | **Frontal Lobe** | | 0 | | | | | |
| L | Superior Frontal Gyrus | Ctx_8Ad_L | 131 | 1048 | -18 | 14 | 48 | Cortex_Default_ModeA |
| L | Inferior Orbitofrontal Cortex | Ctx_a47r_L | 149 | 1192 | -38 | 46 | -12 | Cortex_Fronto_ParietalB |
| L | Middle Orbitofrontal Cortex | Ctx_a47r_L | 124 | 992 | -34 | 54 | -2 | Cortex_Fronto_ParietalB |
| L | Inferior Parietal Lobule | Ctx_PFm_L | 1106 | 8848 | -50 | -50 | 38 | Cortex_Fronto_ParietalB |
| R | Inferior Parietal Lobule | Ctx_PFm_R | 981 | 7848 | 52 | -52 | 40 | Cortex_Fronto_ParietalB |
| R | Middle Frontal Gyrus | Ctx_9_46d_R | 338 | 2704 | 30 | 56 | 22 | Cortex_Ventral_AttentionB |
| L | Middle Frontal Gyrus | Ctx_9_46d_L | 296 | 2368 | -30 | 44 | 26 | Cortex_Ventral_AttentionB |
| R | Middle Frontal Gyrus | Ctx_46_R | 261 | 2088 | 34 | 40 | 34 | Cortex_Ventral_AttentionB |
| L | Middle Frontal Gyrus | Ctx_46_L | 317 | 2536 | -32 | 28 | 36 | Cortex_Ventral_AttentionB |
| | | | | **Non-Modifiable Emotion** | | | | |
| | **Cerebellum** | | | | | | | |
| R | Cerebellum | Multiple regions | 2846 | 22768 | 40 | -68 | -18 | Cerebellum |
| | **Occipital Lobe** | | | | | | | |
| L | Fusiform Gyrus | Multiple regions | 2222 | 17776 | -36 | -68 | -16 | Cortex_Dorsal_AttentionA |
| | **Temporal Lobe** | | | | | | | |
| L | Middle Temporal Gyrus | Ctx_PHT_L | 474 | 3792 | -52 | -60 | 2 | Cortex_Fronto_ParietalA |
| | **Parietal Lobe** | | | | | | | |
| R | Posterior Cingulate Cortex | Ctx_7m_R | 1044 | 8352 | 2 | -46 | 22 | Cortex_Default_ModeA |
| R | SupraMarginal | Ctx_PF_R | 55 | 440 | 62 | -28 | 46 | Cortex_Ventral_AttentionA |
| R | Inferior Parietal Lobule | {'Multiple regions'} | 2702 | 21616 | 34 | -54 | 50 | Cortex_Dorsal_AttentionA |
| L | Inferior Parietal Lobule | {'Multiple regions'} | 2775 | 22200 | -26 | -60 | 48 | Cortex_Dorsal_AttentionB |
| | **Frontal Lobe** | | | | | | | |
| R | Medial Prefrontal Cortex | Ctx_9m_R | 299 | 2392 | 4 | 60 | 24 | Cortex_Default_ModeA |
| L | Medial Prefrontal Cortex | Ctx_d32_L | 97 | 776 | -8 | 44 | 24 | Cortex_Default_ModeA |
| R | Middle Frontal Gyrus | Ctx_6a_R | 1622 | 12976 | 28 | 0 | 56 | Cortex_Dorsal_AttentionB |
| L | Middle Frontal Gyrus | Ctx_6a_L | 1101 | 8808 | -24 | 0 | 56 | Cortex_Dorsal_AttentionB |
| L | Inferior Frontal Gyrus, Opercular | Ctx_44_L | 701 | 5608 | -52 | 10 | 28 | Cortex_Default_ModeB |
| R | Rectus | Ctx_10pp_R | 1062 | 8496 | 0 | 58 | -20 | Cortex_Limbic |
| L | Middle Orbitofrontal Cortex | Ctx_13l_L | 334 | 2672 | -28 | 34 | -20 | Cortex_Limbic |
| R | Inferior Frontal Cortex | Multiple regions | 3746 | 29968 | 40 | 24 | 12 | Cortex_Ventral_AttentionA |
| R | Supplementary Motor Area | Ctx_SCEF_R | 148 | 1184 | 10 | 10 | 48 | Cortex_Ventral_AttentionA |
| L | Inferior Frontal Gyrus, Triangular | Ctx_46_L | 695 | 5560 | -50 | 38 | 18 | Cortex_Ventral_AttentionB |
| R | Middle Cinglate Cortex | Ctx_33pr_R | 448 | 3584 | 0 | 2 | 30 | Cortex_Ventral_AttentionB |

| | | | | | | | | |
|---|---|---|---|---|---|---|---|---|
| R | Middle Orbitofrontal Cortex | Ctx_11l_R | 79 | 632 | 22 | 52 | -18 | Cortex_Fronto_ParietalB |
| | **Subcortical** | | | | | | | |
| R | Thalamus | Thal_LP | 141 | 1128 | 14 | -20 | 12 | Diencephalon |
| R | Thalamus | Thal_VL | 105 | 840 | 6 | -14 | 14 | Diencephalon |
| | Amygdala Parahippocampal region Hippocampus | Multiple regions | 8399 | 67192 | 4 | -8 | -12 | Cortex_Limbic |
| R | Putamen | Putamen_Pa_R | 55 | 440 | 24 | 10 | 4 | Basal_ganglia |
| R | Caudate | Cau_R | 67 | 536 | 6 | 20 | 16 | Basal_ganglia |
| **Modifiable Emotion** | | | | | | | | |
| | **Occipital Lobe** | | | | | | | |
| R | Middle Occipital Gyrus Inferior Occipital Gyrus Superior Parietal Lobule | Multiple regions | 5740 | 45920 | -28 | -68 | 30 | Cortex_Dorsal_AttentionB |
| L | Middle Occipital Gyrus Inferior Occipital Gyrus | Multiple regions | 5501 | 44008 | 34 | -72 | 6 | Cortex_Default_ModeC |
| | **Parietal Lobe** | | | | | | | |
| R | Postcentral | Ctx_2_R | 1404 | 11232 | 54 | -24 | 42 | Cortex_Dorsal_AttentionB |
| L | Precentral | Ctx_6r_L | 334 | 2672 | -52 | 4 | 28 | Cortex_Ventral_AttentionA |
| L | Precuneus | Ctx_v23ab_L | 49 | 392 | -8 | -52 | 16 | Cortex_Default_ModeA |
| R | Precuneus | Ctx_POS1_R | 281 | 2248 | 6 | -52 | 16 | Cortex_Default_ModeC |
| | **Frontal Lobe** | | | | | | | |
| L | Superior Frontal Gyrus | Ctx_6a_L | 150 | 1200 | -24 | -8 | 56 | Cortex_Dorsal_AttentionB |
| R | Inferior Orbitofrontal Cortex | Ctx_11l_R | 95 | 760 | 28 | 34 | -14 | Cortex_Fronto_ParietalB |

Center coordinates of clusters in identified system components are reported in standard MNI space. CanLab_Atlas labels are from HCP-MMP1.0. atlas (Glasser et al., 2016). Cortical network labels are from the 16-network cortical brain parcellation of resting-state fMRI data (Yeo et al., 2011)

**Table S2** Multivariate prediction model using different combination of brain activations in four system components

| Model combination | Coefficent A | Coefficent B | Coefficent C | Coefficent D | Ajusted R square | p value | RMSE | AIC |
|---|---|---|---|---|---|---|---|---|
| A | 0.92 | | | | 0.039 | 0.0002 | 0.647 | 706 |
| B | | 0.64 | | | 0.02 | 0.008 | 0.653 | 713 |
| C | | | 0.11 | | -0.002 | 0.77 | 0.66 | 720 |
| D | | | | -0.82 | 0.017 | 0.008 | 0.653 | 713 |

| | | | | | | | | |
|---|---|---|---|---|---|---|---|---|
| AB | 1.41 | -0.56 | | | 0.037 | 0.0004 | 0.646 | 706.6 |
| AC | 1.42 | | -1.27 | | 0.0516 | 3.02E-05 | 0.641 | 701.1 |
| AD | 1.04 | | | -1.02 | 0.063 | 3.75E-06 | 0.638 | 696.9 |
| BC | | 1.43 | -1.66 | | 0.035 | 0.0007 | 0.647 | 707.4 |
| BD | | 0.95 | | -1.22 | 0.058 | 2.51E-05 | 0.641 | 700.7 |
| CD | | | 1.53 | -1.66 | 0.037 | 4.70E-04 | 0.646 | 706.6 |
| ABC | 1.24 | 0.26 | -1.43 | | 0.05 | 1.00E-04 | 0.642 | 702.9 |
| ABD | 0.98 | 0.07 | | -1.04 | 0.06 | 1.56E-05 | 0.639 | 698.8 |
| ACD | 1.08 | | -0.12 | -0.96 | 0.06 | 1.55E-05 | 0.639 | 698.8 |
| BCD | | 0.98 | -0.094 | -1.18 | 0.05 | 9.71E-05 | 0.642 | 702.7 |
| ABCD | 0.99 | 0.15 | -0.22 | -0.95 | 0.058 | 5.00E-05 | 0.639 | 700.8 |

**A: Reappraisal only**
**B: Common appraisal**
**C: Non-mondifiable Emotion**
**D: Modifiable Emotion**

Using AIC as model selection criteria, combination of ‘Reappraisal only’ and ‘Modifiable emotion’ are the best predictor of reappraisal success.

**Table S3** Neurosynth topic maps and their relative correlations with identified system components.

| *Topic ID in Neurosynth* | *TopicLabel* | *Reappraisal only* | *Common appraisal* | *Non-modifiable emotion* | *Modifiable Emotion* | *PCA1* | *PCA2* | *Selectivity* |
|---|---|---|---|---|---|---|---|---|
| 0 | **Sensory Stimulation** | -0.75 | -1.07 | -1.36 | 0.80 | -1.40 | -1.25 | 1.55 |
| 2 | **Substance Abuse** | 0.21 | -0.35 | -0.56 | -0.36 | 0.15 | -0.73 | 0.56 |
| 4 | **Trauma & Stress** | 0.81 | -0.35 | -0.07 | -0.36 | 0.49 | -0.29 | 0.88 |
| 5 | **Cognitive Impairment** | 0.09 | -0.67 | -0.58 | -0.51 | -0.04 | -0.94 | 0.60 |
| 6 | **Object Recognition** | -1.02 | -1.06 | 0.67 | 4.29 | -3.45 | 1.86 | 3.62 |
| 7 | **Body Cycles** | -0.17 | -0.60 | -0.67 | -0.49 | -0.16 | -1.00 | 0.32 |
| 9 | **Spatial Attention** | -0.59 | 0.09 | 1.98 | 2.01 | -1.47 | 2.46 | 0.02 |
| 13 | **Executive Function** | 1.92 | 0.61 | 0.05 | -0.40 | 1.76 | 0.24 | 1.30 |
| 14 | **Mental Imagery** | -0.26 | 0.54 | 1.78 | 0.70 | -0.33 | 1.94 | 1.08 |
| 15 | **Reading & Writing** | -0.18 | 1.05 | 0.47 | 0.04 | 0.45 | 0.79 | 0.57 |

| | | | | | | | | |
|---|---|---|---|---|---|---|---|---|
| 17 | **Empathy & Interaction** | 1.65 | 0.73 | -0.65 | -0.77 | 1.91 | -0.45 | 0.92 |
| 20 | **Word Processing** | 0.18 | 0.36 | -0.26 | -0.43 | 0.56 | -0.24 | 0.19 |
| 21 | **Contextual Cues** | 0.24 | -0.29 | 0.02 | -0.35 | 0.15 | -0.22 | 0.21 |
| 22 | **Depression & Disorders** | 0.36 | -0.41 | -0.33 | -0.35 | 0.18 | -0.55 | 0.69 |
| 23 | **Creativity & Acupuncture** | -0.13 | -0.55 | -0.04 | -0.42 | -0.20 | -0.43 | 0.08 |
| 24 | **Encoding & Retrieval** | -0.10 | 0.59 | -0.49 | -0.64 | 0.64 | -0.45 | 0.69 |
| 25 | **Response Inhibition** | 5.01 | 1.64 | -0.74 | -0.51 | 4.44 | 0.16 | 3.37 |
| 30 | **Multiple Sclerosis** | 0.29 | -0.27 | -0.50 | -0.33 | 0.23 | -0.63 | 0.57 |
| 32 | **Language Processing** | -0.02 | 1.19 | -0.75 | -0.51 | 1.00 | -0.37 | 1.21 |
| 34 | **Decision-making** | -0.23 | 0.36 | -0.61 | -0.56 | 0.39 | -0.61 | 0.60 |
| 40 | **Action Observation** | -0.81 | -0.72 | 0.18 | 1.94 | -1.91 | 0.62 | 1.76 |
| 46 | **Hearing Disorders** | 0.04 | -0.41 | -0.52 | -0.34 | -0.01 | -0.72 | 0.38 |
| 49 | **Cognitive Conflict** | 0.17 | 2.10 | -0.44 | -0.43 | 1.60 | 0.29 | 1.93 |
| 51 | **Motor Coordination** | -0.78 | -0.51 | -1.01 | -0.36 | -0.53 | -1.23 | 0.15 |
| 52 | **Sleep & Math** | -0.16 | 0.19 | 2.00 | -0.07 | -0.11 | 1.68 | 1.81 |
| 56 | **Memory & Events** | -0.03 | -0.63 | -1.22 | -0.68 | 0.04 | -1.54 | 0.61 |
| 58 | **Task-switching** | 0.78 | 1.97 | 0.07 | -0.39 | 1.85 | 0.71 | 1.18 |
| 60 | **Emotion Processing** | -0.64 | -0.58 | 1.23 | -0.87 | -0.41 | 0.38 | 1.82 |
| 61 | **Pain Perception** | -1.22 | -0.55 | -0.83 | -0.63 | -0.72 | -1.24 | 0.08 |
| 62 | **Developmental Disorders** | 0.28 | -0.27 | -0.34 | -0.34 | 0.21 | -0.51 | 0.54 |
| 65 | **Emotion face** | -0.54 | -0.89 | 2.57 | 0.41 | -1.28 | 1.90 | 2.16 |
| 66 | **Gender Differences** | 0.16 | -0.31 | -0.25 | -0.38 | 0.13 | -0.47 | 0.41 |
| 67 | **Personality Traits** | 0.93 | -0.31 | -0.23 | -0.34 | 0.59 | -0.39 | 1.16 |
| 68 | **Working Memory** | 1.29 | 3.97 | 2.17 | -0.19 | 3.09 | 3.32 | 1.81 |
| 70 | **Body perception** | -0.11 | -0.55 | 0.06 | 1.19 | -0.99 | 0.33 | 1.14 |
| 72 | **Motor control** | -1.11 | -0.96 | -1.30 | -0.03 | -1.15 | -1.53 | 0.93 |
| 75 | **Spatial cognition** | -0.90 | -0.76 | 1.09 | 1.63 | -1.91 | 1.22 | 0.54 |
| 77 | **Reasoning & evaluation** | 0.40 | 0.42 | -0.40 | -0.46 | 0.76 | -0.33 | 0.02 |
| 81 | **Alcohol dependence** | 0.20 | -0.34 | -0.28 | -0.36 | 0.13 | -0.50 | 0.48 |
| 82 | **Task performance** | -0.25 | 1.23 | 1.51 | -0.15 | 0.52 | 1.63 | 0.28 |
| 85 | **Feedback & learning** | 0.15 | 1.21 | -0.46 | -0.50 | 1.10 | -0.11 | 1.06 |
| 86 | **Auditory processing** | -0.69 | -0.92 | -1.46 | -0.79 | -0.47 | -1.93 | 0.10 |
| 87 | **Language comprehension** | -0.16 | 0.86 | -1.22 | -0.75 | 0.88 | -0.99 | 1.03 |
| 88 | **Motion perception** | -0.58 | -0.76 | -0.58 | 2.73 | -2.13 | 0.33 | 3.31 |
| 90 | **Familiarity & recognition** | 0.19 | -0.16 | -0.18 | -0.29 | 0.19 | -0.31 | 0.35 |
| 93 | **Eye movements** | -0.71 | -0.48 | 1.02 | 0.64 | -1.13 | 0.87 | 0.37 |
| 95 | **Motor movements** | -1.73 | -1.45 | -1.15 | 0.73 | -2.22 | -1.32 | 1.88 |
| 97 | **Fear conditioning** | -0.59 | -0.67 | 1.56 | -0.64 | -0.58 | 0.72 | 2.16 |
| 98 | **Food & weight** | -0.28 | -0.60 | 0.56 | -0.50 | -0.33 | 0.00 | 0.83 |
| 99 | **Reward processing** | -0.63 | -0.67 | 0.47 | -0.64 | -0.51 | -0.18 | 1.09 |

PCA was conducted on the correlation matrix and the loadings of each topic maps on first two components are reported. Selectivity is done by the difference between first and second largest correlated coefficients across four system components.

**Table S4** Spatial correlation (std) and meta information for neural transmitter maps. Mean and standard deviation are computed from spatial correlation results with 100 bootstrap samples.

| *target_StudyNum* | *Common appraisal (std)* | *Reappraisal only (std)* | *Non-modifiable emotion (std)* | *Modifiable emotion (std)* | *N* | *age_years* | *primary_reference* |
|---|---|---|---|---|---|---|---|
| 5HT1a_1 | 0.008(0.013) | 0.013(0.015) | -0.035(0.014) | 0.002(0.005) | 8 | 28.4 | beliveau2017jneurosci |
| 5HT1a_2 | 0.061(0.012) | 0.038(0.012) | 0.012(0.014) | 0.033(0.006) | 35 | 26.3 | savli2012neuroimage |
| 5HT1b_1 | 0.128(0.016) | 0.019(0.017) | 0.024(0.013) | 0.087(0.008) | 36 | 27.8 | beliveau2017jneurosci |
| 5HT1b_2 | 0.15(0.016) | 0.058(0.015) | 0.058(0.013) | 0.105(0.01) | 65 | 33.73 | gallezot2010jcerebbloodflowmetab |
| 5HT1b_3 | 0.149(0.015) | 0.048(0.019) | -0.044(0.015) | 0.055(0.008) | 23 | 28.7 | savli2012neuroimage |
| 5HT2a_1 | 0.102(0.014) | 0.057(0.016) | -0.012(0.015) | 0.101(0.008) | 19 | 28.2 | savli2012neuroimage |
| 5HT2a_2 | 0.096(0.017) | 0.041(0.015) | -0.018(0.017) | 0.09(0.008) | 29 | 22.6 | beliveau2017jneurosci |
| 5HT2a_3 | 0.088(0.012) | 0.046(0.012) | -0.009(0.012) | 0.081(0.01) | 3 | 41.14 | talbot2012neuroimage |
| 5HT4 | 0.016(0.021) | 0.013(0.012) | 0.01(0.022) | 0.015(0.003) | 59 | 25.9 | beliveau2017jneurosci |
| 5HT6 | 0.116(0.016) | 0.041(0.012) | 0(0.015) | 0.091(0.009) | 30 | 36.6 | radhakrishnan2018jnuclmed |
| 5HTT_1 | -0.077(0.013) | -0.042(0.006) | 0.155(0.026) | -0.092(0.011) | 18 | 30.5 | savli2012neuroimage |
| 5HTT_2 | -0.057(0.013) | -0.035(0.007) | 0.155(0.025) | -0.048(0.006) | 100 | 25.1 | beliveau2017jneurosci |
| CB1_1 | 0.197(0.018) | 0.097(0.018) | 0.012(0.014) | 0.083(0.012) | 22 | 27.5 | laurikainen2019neuroimage |
| CB1_2 | 0.107(0.012) | 0.079(0.013) | 0.018(0.014) | -0.003(0.006) | 77 | 30.01 | normandin2015jcerebbloodflowmetab |
| D1 | 0.015(0.022) | -0.004(0.008) | 0.025(0.019) | -0.04(0.006) | 13 | 33 | kaller2017eurjnuclmedmolimaging |
| D2_1 | -0.017(0.02) | -0.009(0.006) | 0.043(0.021) | -0.035(0.004) | 49 | 18.41 | jaworska2020neuropsychopharmacology |
| D2_2 | -0.006(0.017) | 0.001(0.007) | 0.066(0.02) | -0.015(0.003) | 37 | 48.36 | smith2017jcerebbloodflowmetab |
| D2_3 | -0.009(0.02) | 0.003(0.007) | 0.066(0.021) | -0.023(0.004) | 55 | 32.45 | sandiego2015jcerebbloodflowmetab |
| D2_4 | 0.016(0.018) | 0.006(0.007) | 0.018(0.02) | 0.023(0.002) | 7 | 24 | alakurtti2015jcerebbloodflowmetab |
| DAT_1 | -0.024(0.018) | -0.015(0.006) | 0.01(0.019) | -0.021(0.003) | 6 | 31.06 | sasaki2012jnuclmed |
| DAT_2 | -0.069(0.021) | -0.042(0.006) | 0.07(0.022) | -0.063(0.01) | 174 | 61 | dukart2018scirep |
| GABAa | 0.12(0.011) | 0.052(0.009) | -0.012(0.013) | 0.101(0.009) | 6 | 43 | dukart2018scirep |
| GABAabz | 0.098(0.011) | 0.036(0.011) | -0.018(0.015) | 0.102(0.011) | 16 | 26.6 | norgaard2021neuroimage |
| H3 | 0.119(0.017) | 0.001(0.008) | 0.062(0.019) | -0.05(0.005) | 8 | 31.69 | gallezot2017jcerebbloodflowmetab |
| mGluR5_1 | 0.1(0.014) | 0.069(0.012) | 0.034(0.011) | 0.09(0.009) | 22 | 67.9 | PI: Pedro Rosa-Neto |
| mGluR5_2 | 0.137(0.016) | 0.058(0.011) | 0.025(0.012) | 0.068(0.008) | 28 | 33.1 | dubois2016eurjnuclmedmolimaging |
| mGluR5_3 | 0.14(0.017) | 0.048(0.012) | 0.016(0.013) | 0.084(0.01) | 73 | 19.9 | smart2019eurjnuclmedmolimaging |
| MOR_1 | 0.114(0.017) | 0.051(0.012) | 0.05(0.019) | -0.131(0.014) | 39 | 39.38 | turtonen2021biolpsychiatrycnni |
| MOR_2 | 0.157(0.02) | 0.067(0.016) | 0.062(0.019) | -0.087(0.014) | 204 | 32.3 | kantonen2020neuroimage |
| NET_1 | -0.009(0.007) | 0(0.006) | 0.067(0.018) | -0.024(0.008) | 10 | 33.3 | hesse2017eurjnuclmedmolimaging |
| NET_2 | 0.037(0.011) | 0.022(0.009) | 0.045(0.013) | 0.089(0.009) | 77 | 33.4 | ding2010synapse |
| VAChT_1 | -0.03(0.018) | -0.017(0.006) | 0.017(0.019) | -0.052(0.005) | 4 | 37 | PI: Lauri Tuominen & Synthia Guimond |
| VAChT_2 | -0.039(0.016) | -0.008(0.007) | 0.017(0.02) | -0.046(0.005) | 5 | 68.3 | bedard2019sleepmed |
| VAChT_3 | -0.015(0.016) | -0.005(0.006) | 0.029(0.019) | -0.036(0.005) | 18 | 66.8 | aghourian2017molpsychiatry |

| | | | | | | | |
|---|---|---|---|---|---|---|---|
| a4b2 | 0.06(0.011) | 0.017(0.01) | 0.044(0.016) | -0.014(0.005) | 30 | 33.5 | hillmer2016neuroimage |
| M1 | 0.108(0.014) | 0.04(0.012) | 0.024(0.014) | 0.111(0.01) | 24 | 40.45 | naganawa2020jnuclmed |
